# Chromatin and gene-regulatory dynamics of the developing human cerebral cortex at single-cell resolution

**DOI:** 10.1101/2020.12.29.424636

**Authors:** Alexandro E. Trevino, Fabian Müller, Jimena Andersen, Laksshman Sundaram, Arwa Kathiria, Anna Shcherbina, Kyle Farh, Howard Y. Chang, Anca M. Paşca, Anshul Kundaje, Sergiu P. Paşca, William J. Greenleaf

## Abstract

Genetic perturbations of cerebral cortical development can lead to neurodevelopmental disease, including autism spectrum disorder (ASD). To identify genomic regions crucial to corticogenesis, we mapped the activity of gene-regulatory elements generating a single-cell atlas of gene expression and chromatin accessibility both independently and jointly. This revealed waves of gene regulation by key transcription factors (TFs) across a nearly continuous differentiation trajectory into glutamatergic neurons, distinguished the expression programs of glial lineages, and identified lineage-determining TFs that exhibited strong correlation between linked gene-regulatory elements and expression levels. These highly connected genes adopted an active chromatin state in early differentiating cells, consistent with lineage commitment. Basepair-resolution neural network models identified strong cell-type specific enrichment of noncoding mutations predicted to be disruptive in a cohort of ASD subjects and identified frequently disrupted TF binding sites. This approach illustrates how cell-type specific mapping can provide insights into the programs governing human development and disease.

## INTRODUCTION

Dynamic changes in the activity of cis-regulatory DNA elements, driven by changes in transcription factor (TF) binding, underlie the complex phenotypic transformations that occur during development (Buenrostro et al., 2018; Stergachis et al., 2013). Single cell methods for probing chromatin accessibility have emerged as a sensitive probe for this activity, and, combined with tools to measure single-cell transcriptomes, have the potential to decipher how combinations of transcription factors drive developmental gene expression programs (Kelsey et al., 2017; Klemm et al., 2019). Quantifying the dynamic activity of regulatory elements also enables the principled inference of the time-point or cell type wherein disease-associated genetic variation may impact a developmental process. For instance, it is still unknown how genetic variants associated with neurodevelopmental disease, such as autism spectrum disorder (ASD), interfere with the genetic programs underlying the development of the human cerebral cortex (Rubenstein, 2011; Zhou et al., 2019).

Corticogenesis is a highly orchestrated and dynamic process that results in the formation of the cerebral cortex, and is characterized by the expansion of apical and basal radial glia (RG) and intermediate progenitors in the ventricular and subventricular zones (VZ, SVZ), the inside-out generation of excitatory glutamatergic neurons, and the differentiation of astrocytes and oligodendrocytes (Greig et al., 2013; Molnár et al., 2019; Silbereis et al., 2016). Cell types derived from outside of the dorsal forebrain, including GABAergic neurons, microglia, and some oligodendrocytes, also migrate and integrate into the cerebral cortex during this period (Wonders and Anderson, 2006). Resolving the gene-regulatory dynamics associated with these diverse developmental trajectories and highly heterogeneous cell states requires investigation of both chromatin and gene expression states at single-cell resolution.

To map the gene regulatory logic of human corticogenesis, we generated single-cell chromatin accessibility and RNA expression profiles from human fetal cortical samples spanning 8 weeks during mid-gestation. The paired maps revealed a class of genes with comparatively large numbers of nearby putative enhancers whose accessibility was strongly predictive of gene expression. These genes with predictive chromatin (GPCs) are frequently TFs, and we observed that their local accessibility precedes lineage-specific gene expression in cycling progenitors. We validated these findings using single cell accessibility and expression profiles derived from the same cell (multiomics). Next, we defined a developmental trajectory for cortical glutamatergic neurons, revealing a continuous progression of TF motif activities associated with neuronal specification and migration. We explored the tendency of certain TF motifs to co-occur along this trajectory and derived a network of key TFs that appear to co-regulate one another. In addition, we characterized the lineage potential of glial progenitors and provided evidence for two transcriptionally and epigenetically distinct astrocyte precursor subtypes. Finally, we trained a deep-learning model to infer base pair-resolved, cell type-specific chromatin accessibility profiles from DNA sequence. These models identified sequence motifs that contribute to cell type-specific accessibility and allowed prediction of the potential impact of genetic variants on the chromatin landscape. The predictions prioritized rare *de novo* noncoding genetic variants associated with ASD, which were enriched in case subjects at levels approaching those seen for deleterious protein-coding mutations. We connected these cell type-specific, high-impact mutations to putative downstream effects on gene expression, demonstrating the ability to map the genetic basis of disease with single cell and single base-resolution at key stages of human cortical development.

## RESULTS

### A single-cell regulatory atlas of the developing human cerebral cortex

To capture cellular heterogeneity in the developing cerebral cortex, we created a gene-regulatory atlas using the Chromium platform (10x Genomics) to generate single-cell ATAC-seq (scATAC) and single-cell RNA-seq (scRNA) libraries from four primary human cortex samples at post-conceptional week (PCW) 16, PCW20, PCW21, and PCW24 (**Figure 1A**). Overall, we obtained 57,868 single-cell transcriptomes and 31,304 single-cell epigenomes after quality control and filtering (**Tables S1–S4, Figure S1**). Consistent with previous studies (Fietz et al., 2010; Hansen et al., 2010; Kang et al., 2011; Pollen et al., 2015; Trevino et al., 2020), immunohistochemical analysis of select tissue samples revealed CTIP2^+^ cells in the cortical plate (CP; **Figures 1B inset 1** and **S2A**) and SOX9^+^ cells in the VZ (**inset 3**), SVZ, and outer SVZ (oSVZ, **inset 2**), as well as GFAP^+^ scaffolding spanning the neocortex at PCW17 and PCW21 (**Figures 1C** and **S2B**). As expected, the proliferation marker KI67 colocalized with both GFAP^+^ cells and with PPP1R17^+^ intermediate progenitor cells (IPCs) in the SVZ and oSVZ (**Figures 1C** and **S2B**).

**Figure 1:**
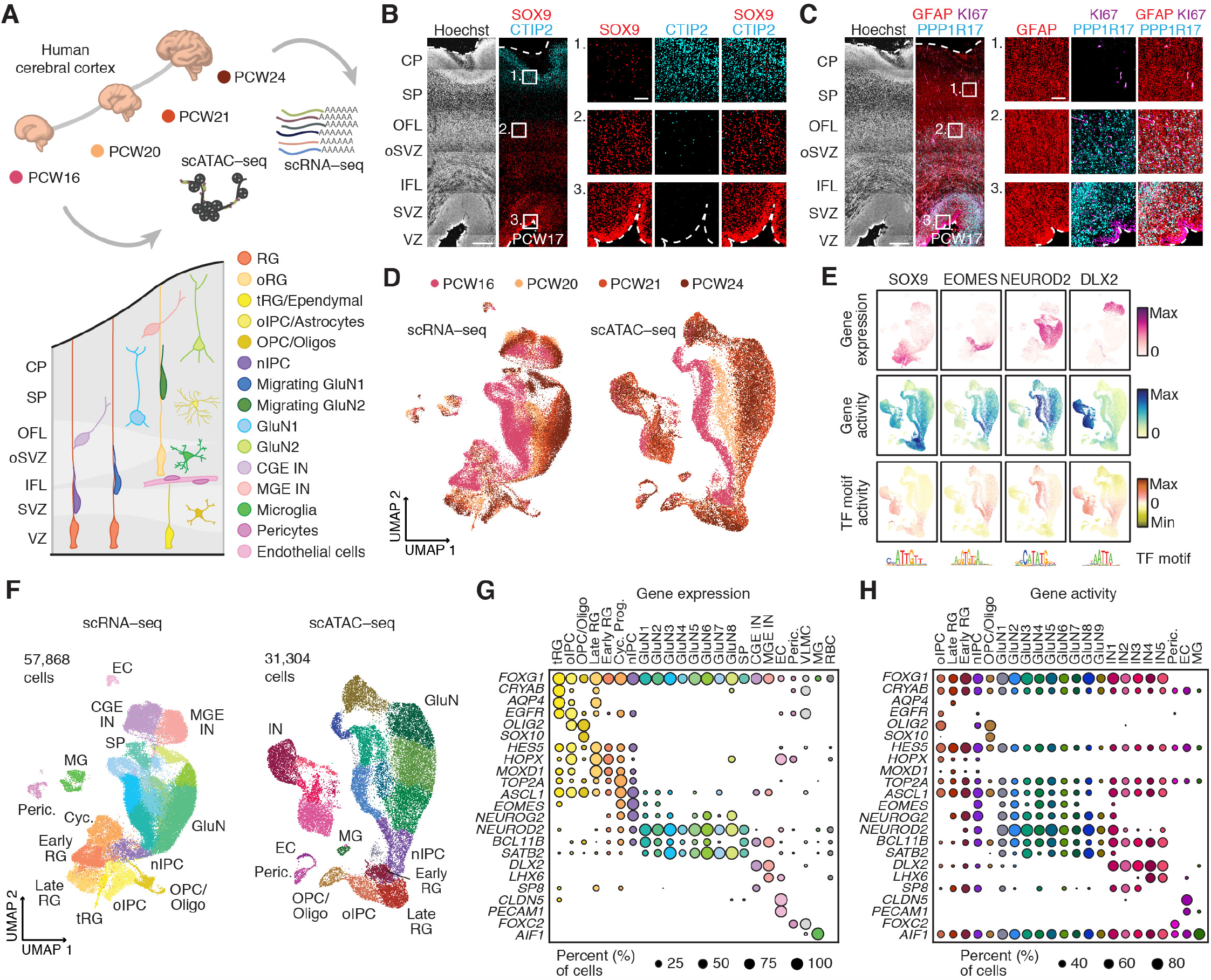
A single cell epigenomic atlas of the developing human neocortex. (A) Schematic of gestational sample time (post-conception week, PCW), genome wide profiling methods and cell types represented in this study. (B) Immunohistochemistry in human cerebral cortex at PCW17 showing expression of SOX9 in VZ, SVZ, and oSVZ, and CTIP2 in cortical plate. Hoechst staining shows nuclei. VZ = ventricular zone, SVZ = subventricular zone, IFL = inner fiber layer, oSVZ = outer SVZ, OFL = outer fiber layer, SP = subplate, CP = cortical plate. (C) Immunohistochemistry in human cerebral cortex at PCW17 showing expression of GFAP, KI67^+^ proliferating cells, and PPP1R17^+^ intermediate progenitor cells. Hoechst staining shows nuclei. (D) Uniform Manifold Approximation and Projection (UMAP) of cells based on gene expression (scRNA-seq, left) and peak accessibility (scATAC-seq, right). Cells are colored according to sample gestational time. (E) Multimodal profiling of SOX9, EOMES, NEUROD2, and DLX2 including gene expression (scRNA-seq), gene activity and TF motif activity (scATAC-seq). (F) UMAP of cells colored by cluster. Cell types labels were assigned based on cluster-specific gene expression and chromatin accessibility. (G) Dot plot showing the percent of cells expressing selected markers across scRNA clusters. (H) Dot plot showing marker gene activity scores derived from chromatin accessibility across scATAC clusters.

To assess global similarities and differences between individual cells, we performed dimension reduction using uniform manifold approximation and projection (UMAP) and clustering. For scATAC, we employed an iterative approach (Granja et al., 2019) to obtain a low-dimensional embedding, cell clustering, and a consensus set of 657,930 accessible peaks representing potential *cis* regulatory elements (CREs; **Methods**). Broadly, the structures of the resulting manifolds for scATAC and scRNA were similar, and they exhibited variation related to gestational time (**Figure 1D**) and cell types (see below). Performing both assays on the same samples enabled us to dissect complementary aspects of gene regulation, including the relationship between gene expression (scRNA) and gene activity (scATAC) – a metric defined by the aggregate local chromatin accessibility of genes (**Methods**) (Pliner et al., 2018), as well as aggregate TF motif activity scores (Schep et al., 2017). Key corticogenesis factors such as *SOX9, EOMES, NEUROD2*, and *DLX2* showed strong cluster-specific enrichments in these three metrics (**Figure 1E**) consistent with their ascribed roles in radial glia (RG), intermediate progenitor cells (IPCs), cortical glutamatergic neurons GluN), and GABAergic neurons (interneuron; IN), respectively.

We next called clusters in both data sets (**Figure 1F**; **Methods**), and annotated these clusters using gene expression and gene activities (**Figures 1G–H** and **S3A, Tables S5–S7, Methods**). In scRNA, we observed a cluster of cycling cells (Cyc) expressing *TOP2A, KI67, CLSPN* and *AURKA*. We also found that radial glia clusters (RG), expressing *SOX9, HES1* and *ATP1A2*, included both ventricular radial glia (vRG: *FBXO32, CTGF, CYR61*) and outer radial glia (oRG: *MOXD1, HOPX, FAM107A, MT3*), and these were separated according to gestational time (early RG, PCW16: *NPY, FGFR3*; late RG, PCW20–24: *CD9, GPX3, TNC*). Cells in one scRNA cluster expressed markers for truncated RG (tRG) and ependymal cells (tRG: *CRYAB, NR4A1, FOXJ1*). In addition to these RG clusters, we identified a cluster expressing genes associated with both RGs and oligodendrocyte lineage precursors (*ASCL1, OLIG2, PDGFRA, EGFR*). This cluster, which we named oligodendrocyte intermediate progenitor cells (oIPC), was different from the oligodendrocyte and oligodendrocyte progenitor cell (OPC/Oligo) cluster that expressed *SOX10, NKX2*.*2* and *MBP*. Astrocytes did not appear to group into a separate cluster, but genes associated with astrocyte identity (*AQP4, APOE, AGT*) were observed in the oIPC cluster and the late RG cluster. A large domain in both representations was composed of neuronal intermediate progenitor cells (nIPC: *EOMES, PPP1R17, PENK, NEUROG1, NEUROG2*), and glutamatergic excitatory neurons (GluN) expressing *NEUROD2, TBR1, BCL11B/CTIP2, SATB2, SLC17A7/VGLUT1*. Among the glutamatergic neuron clusters, we found one group of cells expressing subplate markers (SP: *NR4A2, CRYM, ST18, CDH18*). We also identified distinct clusters of GABAergic interneurons expressing *DLX2, DLX5* and *GAD2*: one of them expressed markers associated with medial ganglionic eminence (MGE)-derived interneurons (MGE: *LHX6, SST*) and the other expressed markers associated with both caudal ganglionic eminence (CGE) and pallial-subpallial boundary (PSB)-derived interneurons (CGE: *SP8, NR2F2*; PSB: *MEIS2, PAX6, ETV1*). In addition, we observed clusters of microglia (MG: *AIF1, CCL3*), and vascular cells including endothelial cells (EC: *CLDN5, PECAM1*), pericytes (Peric: *FOXC2, PDGFRB*), vascular leptomeningeal cells (VLMC: *FOXC2, COL1A1, LUM)*, and red blood cells (RBC: *HEMGN*). Many of the above markers exhibited dynamic gene activity scores in corresponding clusters in scATAC space (**Figure 1H**). While most clusters contained cells representing all gestational time points, some clusters were strongly biased for earlier or later stages (**Figure S3B**). For example, oIPC and tRG clusters were only present in PCW20–24 samples. Projection of another scRNA dataset of the cerebral cortex (Bhaduri et al., 2020) into our scRNA UMAP further corroborated cell type identities and gestational time (**Figure S4**).

We next integrated the derived gene activity scores with gene expression levels, using canonical correlation analysis (CCA) to match cells from one data modality to their nearest neighbors in the other (**Figure 2A**) (Stuart et al., 2019). Cluster annotations of matched cells were consistent across both modalities, except for the cycling progenitor cluster in scRNA, which did not directly map to cells in the chromatin landscape (**Figures 2B** and **S5A, B**). Using pseudobulk aggregates of these matched annotations, we applied a correlation-based approach that links gene-distal CRE accessibility to gene expression (Corces et al., 2018; Ma et al., 2020; Trevino et al., 2020), identifying 64,878 CRE-gene pairs that represent potential regulatory interactions (**Table S8**). Genes in this analysis had a median of 5 linked CREs per gene, with a long-tailed distribution of the number of links. Co-variation of CRE accessibility and gene expression distinguished the identified cell types in both scRNA and scATAC (**Figure 2C**). Clustering of the associated CRE accessibility revealed particularly high variability across clusters corresponding to glial cell populations, corroborated the distinctiveness of GABAergic neuron clusters, and indicated dynamic patterns of gene regulation across glutamatergic neuron clusters.

**Figure 2:**
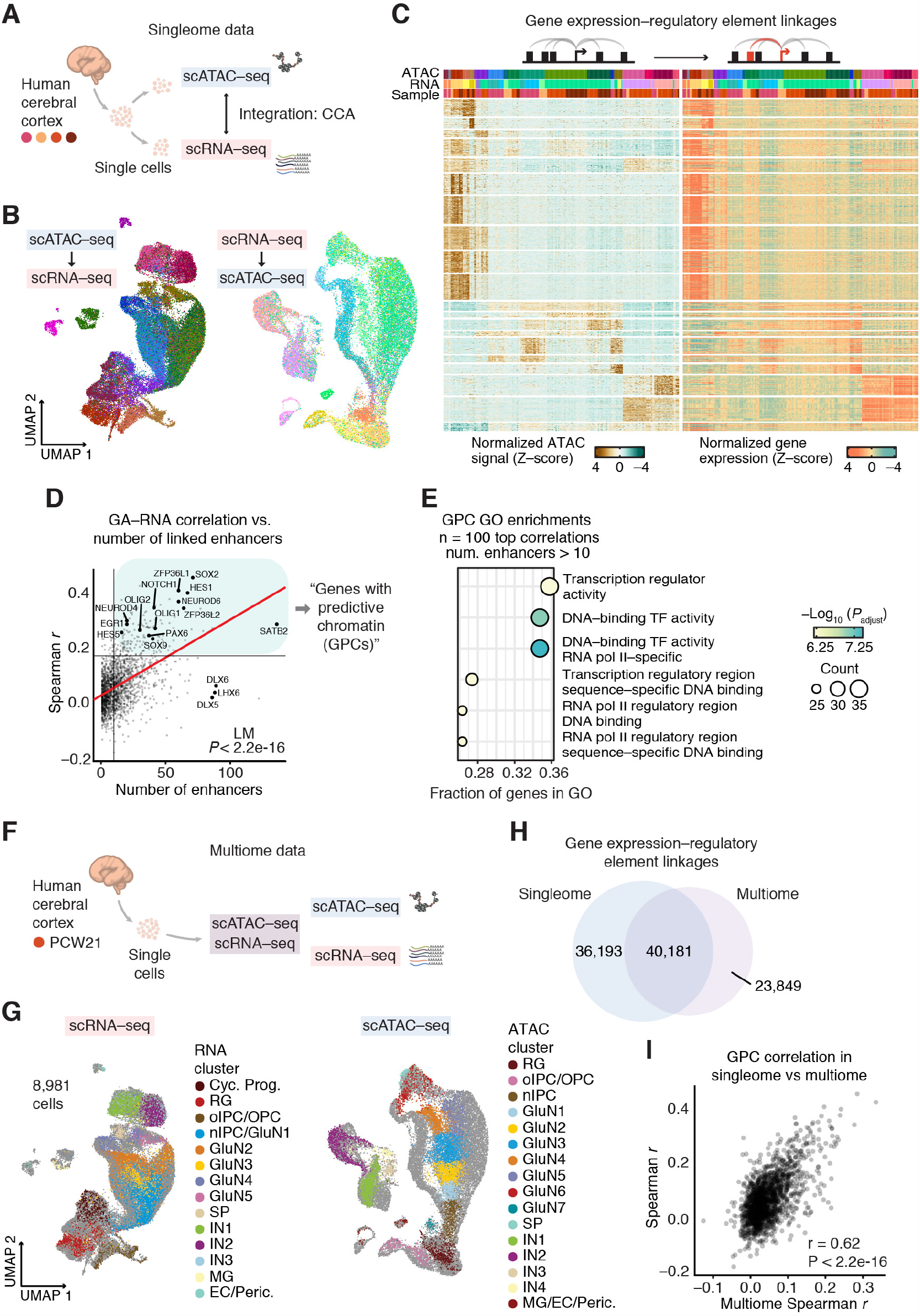
Integrative and multiomic gene regulatory dynamics in the developing human cerebral cortex. (A) Schematic showing the generation and integration of singleome scATAC-seq and scRNA-seq data. Matching is performed by mapping cells into a low-dimensional space using Canonical Correlation Analysis (CCA) and finding nearest-neighbors in that space. (B) UMAPs of scRNA and scATAC cells colored by cluster assignment of matched cells in the respective complementary data modality. (C) Heatmap showing chromatin accessibility and gene expression of 64,878 significantly linked CRE-gene pairs (rows, left CRE accessibility, right linked gene expression) across 200 pseudobulk samples (Methods). Rows were clustered using k-means clustering (k=20). For visualization, 10,000 rows were randomly sampled. (D) Scatterplot showing the correlation between single-cell gene expression and chromatin-derived gene activity (GA), and the number of linked CREs per gene. Transcription factors are labeled. (E) Gene Ontology (GO) enrichment analysis of the 185 genes with predictive chromatin (GPCs) identified in D. (F) Schematic showing the generation of scATAC-seq and scRNA-seq data from the same cells (multiome data) in human cerebral cortex. (G) Projection of multiome scATAC into singleome scATAC UMAP space, and multiome scRNA into singleome scRNA UMAP space. (H) Venn diagram showing overlap of CRE-gene linkages identified in singleome versus multiome data. (I) Correlation scatterplot showing the correspondence between predictive chromatin in singleome versus multiome data. Pearson r = 0.62, *P* < 2.2e-16.

We then asked if there were genes whose expression could be well-predicted from chromatin accessibility signals by ranking single-cell gene activity-expression correlations for each gene. Unsurprisingly, given the relative sparsity of single-cell ATAC-seq and RNA-seq data, few genes exhibited high correlations by this metric (**Figure 2D**). However, the most robustly correlated genes included factors with central roles in corticogenesis, such as *SOX2* and *HES1*, and these genes were linked to greater numbers of putative enhancers (*P* < 2.2e-16). We hypothesized that these comprised a class of highly regulated genes that play a driving role in establishing cell identities in the developing cerebral cortex. Therefore, we defined a set of 185 genes with predictive chromatin (GPCs), which were in the top decile of gene activity-expression correlations and were linked to a minimum of 10 CREs (**Table S9, Figure 2D**). In this gene set, gene ontology (GO) enrichment analysis revealed a strong enrichment of transcription regulator activity and DNA-binding TF activity (**Figure 2E**).

To validate these inferences, we generated joint scATAC and scRNA data in the PCW21 human cerebral cortex (multiome) (**Figure 2F**). Filtering across both data modalities resulted in 8,981 cells with high-quality transcriptome and epigenome profiles (**Tables S10–12, Figure S6**). We projected these multiomic scATAC and scRNA profiles into the corresponding individually generated landscapes and confirmed that our cell type annotations were well represented in the joint data (**Figure 2G**). When we applied our CRE-gene linking approach to the true cell-to-cell matches, we found that 40,181 inferred peak-gene linkages (53%) were validated from this single timepoint measurement, and an additional 23,849 were identified (**Figure 2H, Table S13**). Thus, the majority of inferred CRE-gene interactions were observed when accessibility and expression measurements were made in the same individual single cells. The multiome data allowed us to validate our set of GPCs, and we found a strong concordance of gene activity-expression correlations between separate cells linked in ATAC-seq and RNA-seq by our analysis and correlations observed when RNA-seq and ATAC-seq are generated from the same cell (Pearson r = 0.62, *P* < 2.2e-16; **Figure 2I**). Therefore, GPCs are also readily apparent in this joint data set, underlining the correspondence between their local accessibility and their transcription within the same cell.

### Continuous trajectories of gene regulation across cortical neuron differentiation

Glutamatergic projection neurons comprise ∼80% of neurons in the cerebral cortex, and distinct subtypes are born in a specific sequence during development. Although several key factors controlling cell fate in corticogenesis have been described (Greig et al., 2013), the gene-regulatory logic that governs specification, migration, and maturation of neural cells has not yet been resolved in human development. Our paired single-cell atlas provided an opportunity to infer the dynamics of these molecular processes in an unbiased fashion. We therefore focused our analysis on glutamatergic neuron clusters, first annotating each cell with a developmental pseudotime, which was inferred by anchoring a differentiation starting point in the Cyc cluster and applying an algorithm based on diffusion through cell-similarity networks derived from RNA velocities (Bergen et al., 2020; La Manno et al., 2018) (**Figures 3A** and **S7A–D**). To test how the architecture of the adult cerebral cortex mapped onto this trajectory, we projected an independent scRNA-seq data comprising neurons from human cerebral cortex (Hodge et al., 2019) into the developmental landscape and identified the nearest neighbor cell for each adult scRNA-seq profile (**Figure S7E**). Adult glutamatergic neurons projected almost exclusively into the neighborhoods of developmental cells annotated with later pseudotimes (**Figure S7F**). As expected, we observed association of earlier and later gestational timepoints with deeper and upper adult cortical layers respectively (**Figure S7G**). When we compared the expression levels in migrating neurons from the early gestational timepoint (PCW16) to those from later timepoints (PCW20 to PCW24), we observed increased expression of *LIMCH1, RUNX1, SNCB* and *DOK5* and decreased expression of the AP-1 TF family (*JUN, FOS*), heat shock factors *HSPA1A/B* and *DUSP1* (**Figures S7H** and **S7I, Table S14**). Overall, we found surprisingly few differentially expressed genes in this analysis that have been previously implicated in neurogenesis, suggesting that a considerable degree of gene expression and regulatory variability could be associated with pseudotime, rather than gestational time. We therefore decided to investigate the regulatory dynamics along the pseudotime axis.

**Figure 3:**
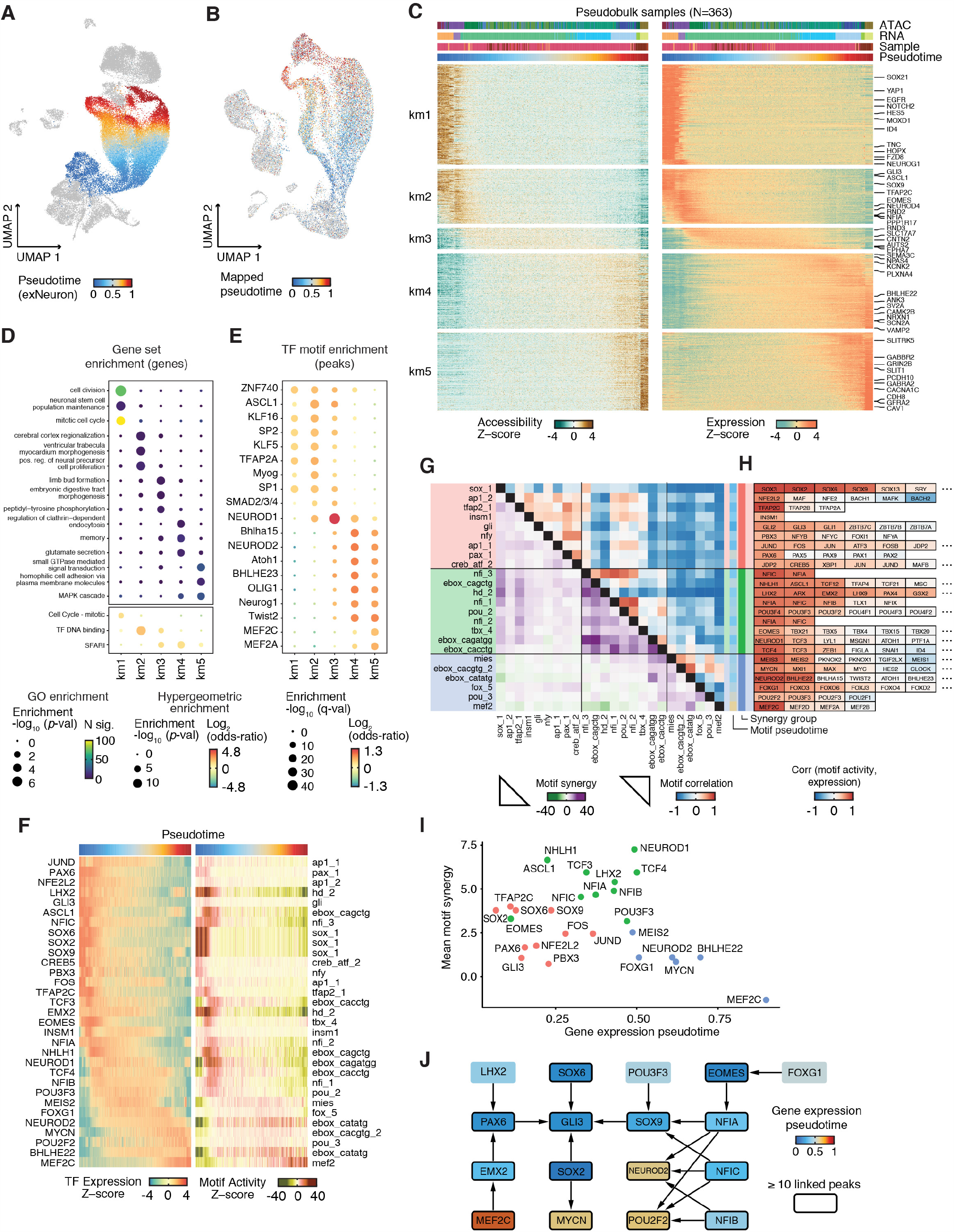
Molecular signatures of excitatory projection neuron generation, migration, and maturation. (A) UMAP of scRNA cells highlighting the glutamatergic neuron trajectory and pseudotime. (B) scATAC UMAP with transferred pseudotime annotation. (C) Heatmap showing accessibility and expression of 13,989 linked CRE-gene pairs (rows, left CRE accessibility, right linked expression) across 363 pseudobulk samples. Interactions (rows) were clustered using k-means clustering (k=5). (D) Gene set enrichment analysis of interaction clusters. Gene ontology (top) and hypergeometric test (bottom) *P*-values are shown along with the number of matched genes or the enrichment odds ratio. (E) Enrichment of TF motifs in peaks represented in interaction clusters. Color represents the odds ratio and size represents the −log10 (*P-*value). (F) Heatmaps showing z-score normalized expression (left) and motif activity (right) of TFs in pseudobulk aggregates. Shown are 31 dynamic TFs associated with 24 motif clusters (Methods). (G) TF motif correlation coefficients (upper triangular heatmap) and synergy z-scores (lower triangular heatmap) of motif clusters in F. Scores were computed using chromVAR. Motifs were hierarchically clustered on synergy z-scores, and this dendrogram was cut to obtain three clusters. (H) Correlation coefficients of TF motif cluster activity and expression. Cluster lists were truncated to the top 6 best correlated genes. (I) Scatterplot showing aggregate gene expression pseudotime versus mean motif synergy. Point colors denote the cluster assignments in G. (J) Network of inferred regulatory interactions (Methods) between TFs in F. Network nodes are colored according to expression weighted pseudotime.

To connect expression trajectories to the accessibility dynamics of specific regulatory elements, we transferred pseudotime values from RNA cells to their nearest ATAC cell neighbors, confirming that this produced a smooth continuum of pseudotime in the chromatin manifold (**Figure 3B**). By applying our correlation-based peak-to-gene linking approach to the glutamatergic neuronal lineage, we identified 13,989 dynamic interactions across pseudotime and grouped these interactions into five clusters (**Figure 3C, Table S15**). Linked genes active early in pseudotime exhibited GO enrichments for cell division and neural precursor proliferation, whereas later interactions were associated with morphogenesis, cell migration and maturation (**Figure 3D**). Interestingly, genes encoding TFs and DNA-binding proteins were particularly enriched in intermediate interactions, while genes from the SFARI database (Abrahams et al., 2013) were more likely to be linked later in pseudotime.

To nominate TFs that may control these dynamic expression programs, we identified TF motifs that were enriched in the different clusters of linked regulatory elements. Motifs enriched in interactions early in the trajectory included ZNF740, KLF16, SP1/2, and ASCL1 (**Figure 3E**). Conversely, interaction clusters associated with intermediate and late pseudotime were associated with motifs of neuronal TFs (NEUROD1/2, NEUROG1, MEF2C). These enrichments represent the putative regulatory vocabulary of individual CREs and their target genes. To characterize the TF-driven regulatory dynamics of neurogenesis over pseudotime in more detail, we linked specific TF genes to TF motifs by correlating TF expression with chromVAR-derived TF motif activity scores. To avoid correlation biases between similar putative binding motifs, we assigned variable TFs to 24 previously defined clusters of motifs (Vierstra et al., 2020) (**Figure 3F**). We observed synchronized TF expression and motif activity for dynamic regulators along neuronal developmental pseudotime, starting with PAX6, SOX2/6/9, GLI3 and ASCL1 motifs, followed by intermediate stage factor motifs (EOMES, NFIA, NFIB, NEUROD1), and finally late-stage motifs (NEUROD2, BHLHE22, MEF2C). Together, these data describe cohesive, sequential waves of motif activations during human corticogenesis that are consistent across gestational time points.

To better understand how TFs are coordinated during human corticogenesis, we next computed the genome-wide synergy and correlation patterns of motif family accessibility (**Figures 3G** and **3H**; **Methods**) (Schep et al., 2017). We found three broad classes of motifs associated with accessibility and TF expression over pseudotime (**Figure 3G–I**): (i) early activity motifs exhibiting moderate synergies (SOX, GLI, PAX) (ii) intermediate activity motifs (NFI/TBX/EOMES) that are highly synergetic within their class, and (iii) late activity motifs that are less cooperative and generally appear to operate more independently (NEUROD2/BHLHE22, MEF2). These findings are broadly consistent with a higher degree of TF motif coordination early in neurogenesis and regulation of later neuronal maturation by a smaller set of more independent TFs. Finally, we derived a TF regulatory network by linking factor-specific motif activity in regulatory elements to TF gene expression (**Figure 3J**; **Methods**). This network indicates that key factors of neurogenesis such as PAX6, SOX2, EOMES and NFIA could regulate effector TFs like NEUROD2, POU2F2 and GLI3 thereby driving later neuronal differentiation, maturation and migration.

### Clustering approach to link gene expression programs to cell fate decisions

We observed extensive heterogeneity in glial cell populations, corresponding to distinct yet partially overlapping expression programs in the identified cell clusters (**Figures S8A** and **S8B**). To develop a high-resolution map of glial populations, we adopted an analysis to identify modules of co-expressed genes. We generated pseudobulk data sets from a k-nearest neighbor (KNN) graph of glial cells, then performed fuzzy c-means clustering on the most variable genes to fractionally assign genes to modules (**Figures 4A** and **S8C** left; **Table S16**). This approach allowed for cells to be annotated with module activities, and for genes to be shared between multiple modules (**Figures S8C** and **S8D**; **Tables S17** and **S18**). This enabled us to explore the relationships between modules and to explore how cells may progress from one module to another across differentiation. To visualize these relationships, we further embedded these cell loadings into a low-dimensional representation of the differentiation landscape (**Figure 4A**, bottom). The structure of this embedding and the underlying module assignments was stable to fuzzy clustering parameters (**Methods**).

**Figure 4:**
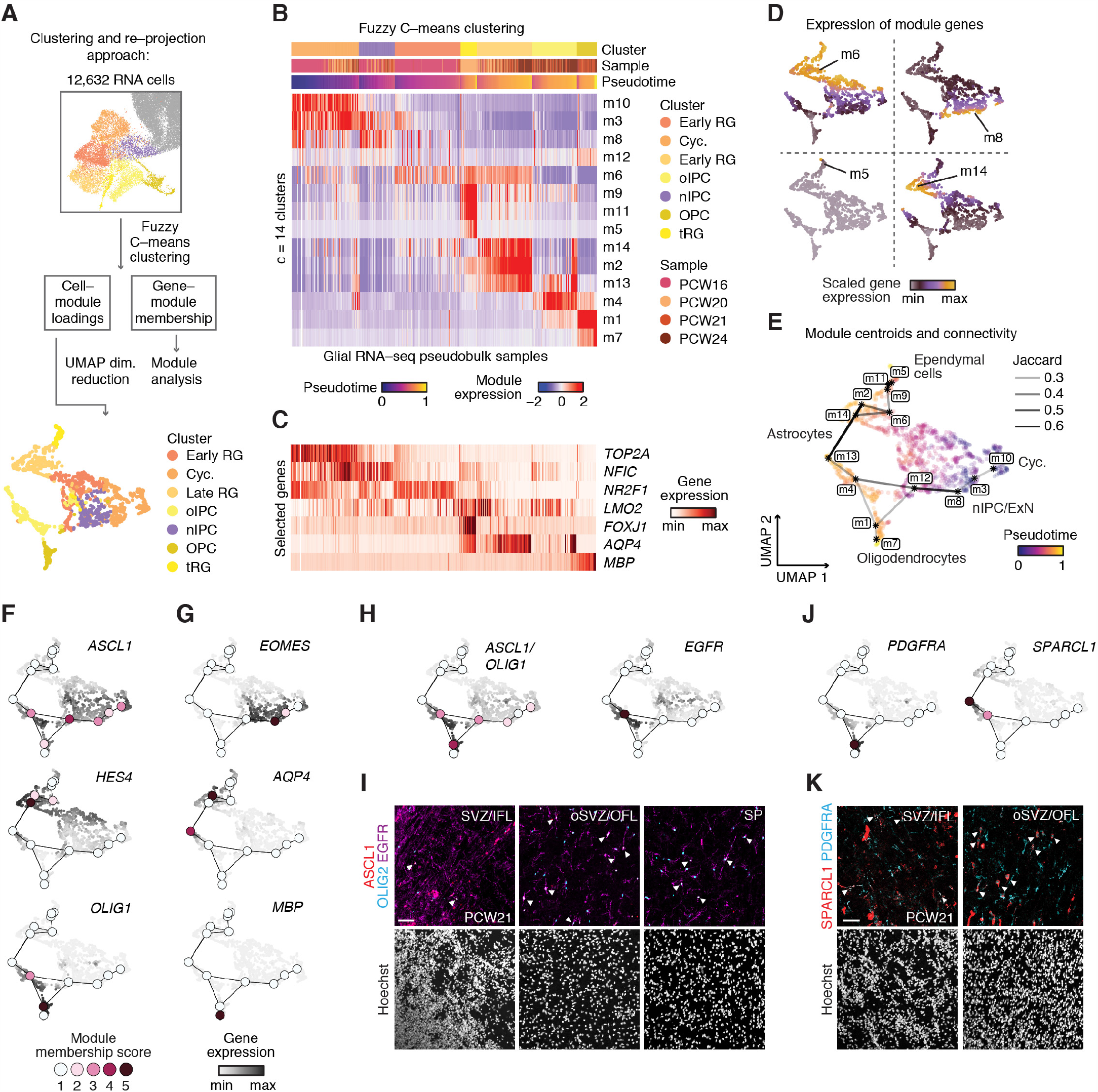
Regulatory logic of glial cell specification. (A) Schematic illustrating the approach used for clustering and reprojection of glial cells by their gene expression. Points in the bott om panel correspond to pseudobulk aggregates of 50 cells. (B) Heatmap of module expression across pseudobulk aggregates, showing variation by cluster, sample age, and pseudotime. (C) Heatmap showing the expression of selected genes across the same pseudobulks. (D) The mean scaled expression of selected gene modules is shown in the low-dimensional UMAP embedding. Figure S9B shows all modules. (E) Projection of module centroids into UMAP space. Pseudobulk samples are colored by pseudotime. Module overlap is shown by lines between centroids and was computed by thresholding the pairwise Jaccard index at > 0.2. (F) Module membership and expression values for factors associated with the three main differentiation programs observed through the glial modules. Module membership scores denote the respective gene’s quantile of membership after zero values are excluded. (G) As in F, for ASCL1, HES4 and OLIG1, factors associated with neuronal intermediate progenitors, astrocytes and oligodendrocytes, which appear as endpoints in this clustering approach. (H) As in F for genes associated with the oIPC cluster of cells and modules m12, m4 and m1. (I) IHC of PCW21 human cerebral cortex showing expression and colocalization (white arrowheads) of ASCL1, OLIG2, and EGFR in cells of the SVZ, oSVZ, outer and inner fiber layers (OFL, IFL) and SP. (J) Module membership and expression values for PDGFRA and SPARCL1, associated with modules m4 and m1, respectively. (K) IHC of PCW21 human cerebral cortex showing expression and colocalization (white arrowheads) of SPARCL1 and PDGFRA in cells of the SVZ, oSVZ and outer and inner fiber layers (OFL, IFL).

To understand the biological basis of these modules, we first examined their expression across cell clusters, developmental stage, and pseudotime (**Figure 4B**), which was rooted in cycling (“Cyc”) cells and correlated with developmental time (Pearson r = 0.67, *P* < 2e-16; **Figure S8E**; **Methods**). Glial maturation genes *FOXJ1, AQP4*, and *MBP*, which are markers for ciliated ependymal cells, astroglia, and oligodendrocytes, respectively (Barbarese et al., 1988; Jacquet et al., 2009; Zhang et al., 2016), were expressed in late-pseudotime cells and assigned primarily to modules m5, m2, and m7. In contrast, the expression of genes associated with cell division and progenitor states, such as *TOP2A, NR2F1*, and *NFIC*, peaked early in pseudotime and were assigned primarily to modules m10, m6, and m3 (**Figures 4C** and **S9A**). Some modules spanned many pseudobulk samples and developmental ages, such as m6 and m8, indicative of sustained longitudinal expression programs, while others were restricted to a few samples or stages, like m5 and m14 (**Figures 4B, 4D** and **S9B**). Modules exhibited distinct GO enrichments, including “cation and metal ion binding” in m6, which may be related to the role of human astrocytes in metal ion homeostasis (Vasile et al., 2017; Zhang et al., 2016), and disease associations (**Figures S9C** and **S9D**). Module m5, comprising *FOXJ1*^+^ cells, was enriched for dynein binding and microtubule activity, consistent with the role of ependymal cilia in circulating the cerebrospinal fluid (Ransom, 2012). When we assessed the expression of some of the genes found in these modules by immunohistochemistry, we found that the transcription factor TFAP2C, which associated with module m6, was expressed in progenitors in the VZ and SVZ (**Figures S10A** and **S10B**). Similarly, PBXIP1, which was associated with m2, was expressed in radial glia in the VZ and SVZ, but not in more mature astrocytes in the CP (**Figures S10C** and **S10D**). CRYAB, associated with m9, was expressed in tRG in the VZ, as previously described (**Figures S10E** and **S10F**) (Nowakowski et al., 2016).

Our clustering and reprojection approach enabled us to compute the degree of gene overlap between modules, which provided a measure of module similarity across our glial landscape (**Figure S10G**). To visualize these relationships, we computed the weighted average of module gene expression across pseudobulk aggregates and plotted these “module centroids” and their connectivity (Jaccard index > 0.2) in the embedding, along with pseudobulks and their pseudotime values (**Figure 4E**). Investigation of module memberships in this representation revealed three broad programs emanating from the cycling cluster: (1) an *ASCL1*^+^ program associated with m3 and m8 and terminating in *EOMES*^+^ nIPCs, (2) a *HES4*^+^ program associated with module m6 and terminating in astrocytes and ependymal cells, and (3) an *ASCL1*^+^/*OLIG1*^+^ program associated with m12, m1, and m4, branching into two endpoints (**Figures 4F** and **4G**). The *ASCL1*^+^/*OLIG1*^+^ program was of particular interest, as it corresponded to the oIPC cluster of cells, which expressed markers associated with both astroglia (*GFAP, HOPX, EGFR, ASCL1*) and oligodendrocyte progenitors (*OLIG2, PDGFRA*), suggestive of a common multipotent glial progenitor (**Figures 4H** and **4J**). To validate the presence of these cells *in situ*, we performed immunohistochemistry for ASCL1, OLIG2 and EGFR in PCW21 cerebral cortex (**Figures 4I, S11** and **S12**). We found that these proteins were often colocalized in the SVZ/IFL, oSVZ/OFL and SP. Next, we reasoned that, if generated from a common glial progenitor, astrocyte and oligodendrocyte precursors might also share expression of markers associated with more differentiated states. To test this, we performed immunohistochemistry for PDGFRA and OLIG2, markers associated with oligodendrocyte progenitors, and SPARCL1, which is a marker associated with mature astrocyte identity (Zhang et al., 2016) (**Figures 4K** and **S13**), and found that they indeed also colocalized in the SVZ/IFL and oSVZ/OFL. We speculate that a subpopulation representing a common multipotent glial progenitor, competent to differentiate into both astrocytes and dorsal forebrain-derived oligodendrocytes, could explain this substantial overlap of expression programs.

### Chromatin and gene expression profiles identify two astrocyte precursor populations

Human cortical astrocytes are larger, more morphologically complex (Oberheim et al., 2009; Zhang et al., 2016), and likely more diverse than those of other mammals (Vasile et al., 2017). However, the developmental steps underlying the diversification of human astrocytes are unknown. We observed three interconnected fuzzy gene modules, largely derived from PCW24 tissue, expressing *AQP4, TNC, ALDH2*, and *APOE*, and other genes specifically expressed in astrocytes (m2, m13, m14) (Sloan et al., 2017; Wiese et al., 2012; Zhang et al., 2016) (**Figures 5A, S14A** and **S14B**). To test whether these transcriptionally related yet distinct subpopulations are associated with different regulatory factors, we computed differential motif enrichments between enhancers linked to genes in two of the modules: m13 versus m14. We found that the bHLH factor motifs ASCL1 and NHLH1 were enriched in module m13, while SOX21 was enriched in m14 (**Figure 5B**). In our glial cells, the accessibility of ASCL1 and NHLH1 motifs correlated best with the gene expression of bHLH factor *OLIG1* (Spearman rho = 0.34 and 0.36, respectively), and we have previously nominated SOX21 as a potential regulator of astrocyte maturation in long-term cortical organoid cultures (Trevino et al., 2020). Thus, two distinct astrocyte-like expression patterns could be distinguished by the chromatin accessibility of *OLIG1* versus *SOX21* motifs.

**Figure 5:**
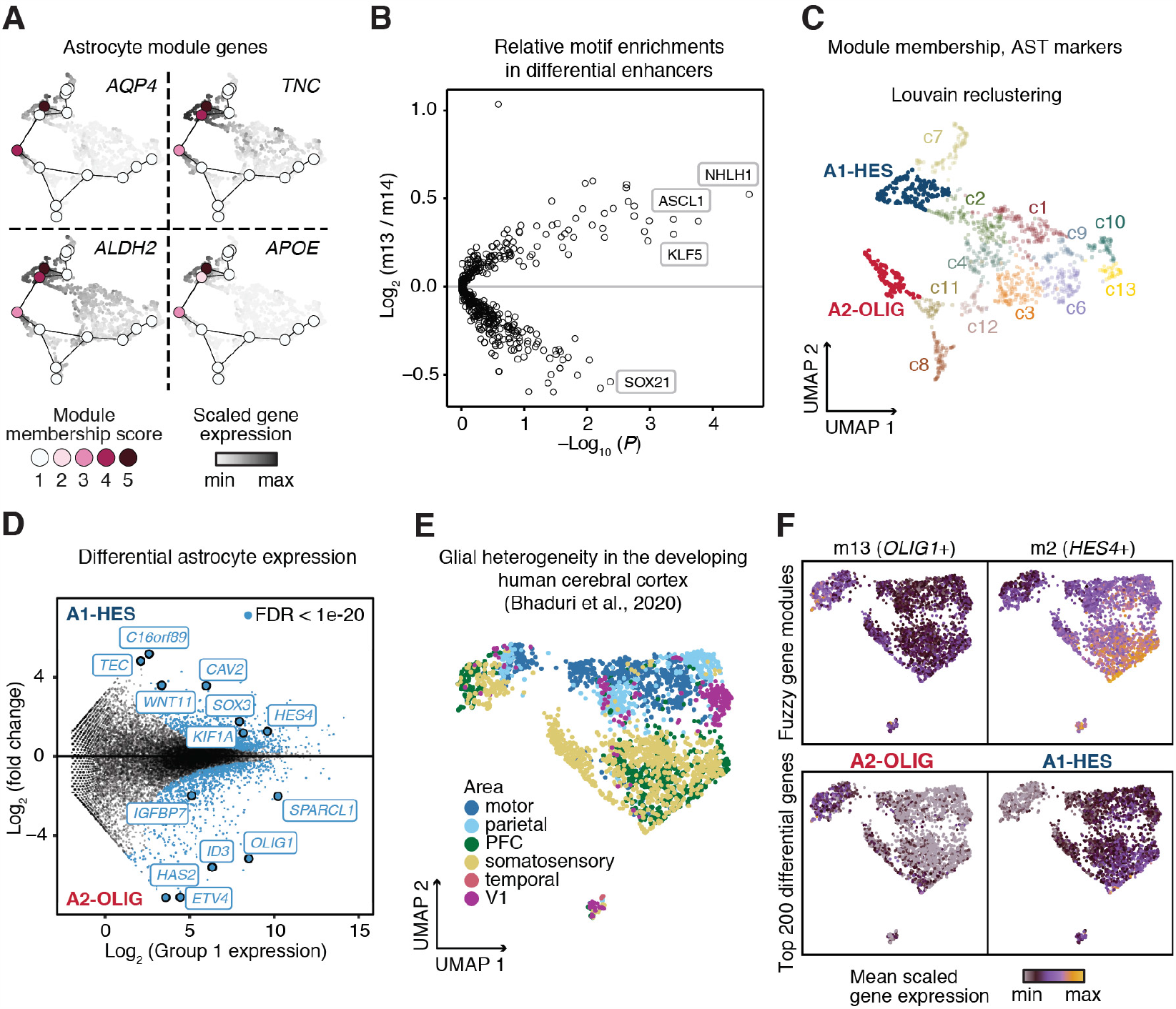
Astrocyte precursor heterogeneity. (A) Module membership and scaled gene expression of astrocyte-associated genes AQP4, TNC, ALDH2 and APOE showing that modules m2, m13 and m14 connect astrocytes. (B) Motif enrichments in peaks linked to module 13 genes relative to peaks linked to module 14. (C) Re-clustering of samples in fuzzy clustering embedding. AQP4 positive clusters are highlighted and defined as A1-HES and A2-OLIG. (D) Differential gene expression between A1-HES and A2-OLIG clusters, calculated using DESeq2. A threshold of Benjamini-Hochberg corrected FDR of 1e-20 was used for visualization (blue). (E) Reanalysis of an orthogonal human fetal scRNA-seq dataset (Bhaduri et al., 2020). Shown is a UMAP of astrocytes colored by cortical area. (F) Mean scaled expression of modules m13 and m2 in orthogonal data, showing partition of module expression into unbiased divisions in the astrocyte UMAP (top), and of the top 200 differential genes from D (bottom).

To examine the differences between cells expressing these modules in more detail, we clustered pseudobulk aggregates to compare cell subsets, and computed differential gene expression between the astrocytic cell clusters A1-HES and A2-OLIG, corresponding to expression of modules m2/14 and m13, respectively (**Figures 5C** and **5D**; **Table S19**). Cluster A1-HES exhibited significantly higher expression of *HES4* and *CAV2*, while A2-OLIG was characterized by increased *SPARCL1, ID3*, and *IGFBP7* expression (**Figures 5D** and **S14C**). To determine if these distinct astrocyte precursor subtypes were due to the sampling of different cortical areas, we used an independent, previously published scRNA-seq dataset of the developing human cortex (Bhaduri et al., 2020) to generate a low-dimensional representation of astroglia (**Figures 5E** and **S14D**). Using this dataset, we visualized the expression of genes identified in our analysis, either from astrocytic modules (m13, m14) or by taking the top 200 most differentially expressed genes from glial cell populations. We found that these gene sets were expressed in distinct cell populations in the independent data set and that this different was not explained by differences in cortical area (**Figure 5F**).

### Chromatin state links GPCs to lineage determination in cycling cells

We next examined how the chromatin state of progenitor cells could potentially affect the acquisition of expression programs characteristic of more differentiated cell states. We therefore focused on the heterogeneity among cells that expressed gene modules strongly associated with cell cycle signatures (**Figure 6A**; Pearson r = 0.89, 0.91 respectively). To link chromatin accessibility to the glial-centric expression map, we generated pseudobulk data sets by sampling local neighborhoods (50-cells) from 13,378 glial scATAC cells. We projected these ATAC-seq pseudo-bulk samples into our gene expression module-derived manifold using accessibility-derived gene activity scores. Consistent with our CCA cluster matching analysis (**Figures 2B** and **S5**), pseudobulks comprised mainly of cells from ATAC cluster c15 (OPC/Oligo) projected into the oligodendrocyte endpoint of this map; cluster c10 (oIPC) data projected into the ASCL1^+^/OLIG2^+^ astrocyte compartment; and cluster c9 (late RG) data projected into both ependymal and HES4^+^ astrocyte endpoints (**Figure 6B**). However, while no distinct cycling cluster was formed in the independent ATAC-seq clustering, a subset of these ATAC-seq pseudobulk samples projected into the cycling, early-pseudotime compartment of the RNA-seq embedding. These samples partitioned into three distinct branches defined by their scATAC cluster assignments (**Figure 6C**; branches A, B, and C). We speculate that strong cell cycle signatures in RNA-seq may have diminished these distinctions that are more clearly seen in ATAC-seq data, and that analyzing these separate branches might allow us to determine if cycling progenitors are poised towards distinct post-mitotic fates, and what factors influence these fate decisions.

**Figure 6:**
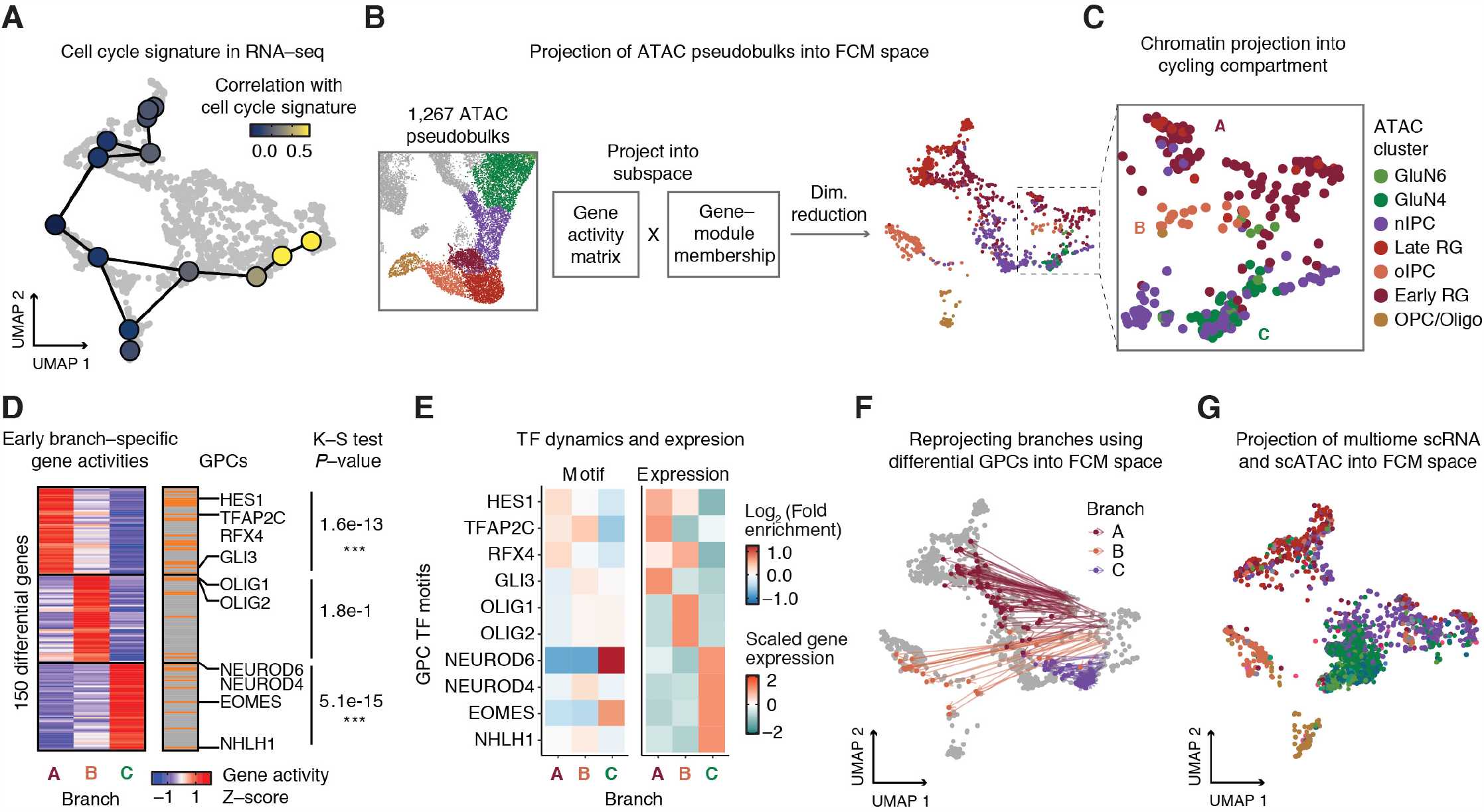
Chromatin state links GPCs to cell fates. (A) Pearson correlation of a cell cycle signature (MSigDB) with module expression signature across pseudobulks. (B) Schematic of ATAC-seq projection into fuzzy clustering embedding. (C) Projection of ATAC-seq pseudobulks into Cyc cluster, and in the neighborhood of cycling-associated modules. Branches are defined. (D) Heatmap showing the 50 most uniquely active genes in branches A, B and C. Gene activities are row-scaled. GPCs are shown to the right of heatmap as orange bars. Select GPCs, which are also TFs, are highlighted. The P-value of a Kolmogorov-Smirnov test for enrichment of GPCs in differential, branch-specific genes is shown, *** indicates that P < 1e-10. (E) Dynamics of GPC motifs and gene expression across three branches of cycling cells. Heatmaps represent enrichment of GPC TF motifs (left) and gene expression levels (right) in branch aggregates. (F) Reprojection of branch A, B, and C using only chromatin accessibility associated with GPCs, showing specific alignment into more mature states. (G) Projection of multiome scRNA data into fuzzy clustering embedding. Cells (points) are colored by the corresponding multiome scATAC cluster.

To explore factors that influence these fate decisions, we identified the 50 most unique genes for each branch based on their gene activity scores (**Methods**). Strikingly, we observed a strong overlap of these genes with the set of GPCs, including *HES1, RFX4, OLIG1, OLIG2, NEUROD6*, and *EOMES*. Overall, differential chromatin activity in all three branches of cycling cells was enriched for GPCs (Kolmogorov-Smirnov test, P = 1.6e-13, 1.8e-1, and 5.1e-15, respectively; **Figure 6D**). For TF GPCs, we computed target motif enrichments across branches, as well as matching gene expression values (**Figure 6E**). Each branch contained at least one basic helix-loop-helix (bHLH) GPC TF in the top 5 most unique genes (*BHLHE40, OLIG1, OLIG2, NEUROD6, NEUROD4*). The similarity of annotated motifs for these factors is consistent with the hypothesis that they can compete for similar binding sites to drive multiple distinct cell fates, as has been previously suggested (Imayoshi et al., 2013; Zhou and Anderson, 2002). Together, these results suggest that differential chromatin activity as well as gene expression of GPCs appear to be prominent features that distinguish different types of cycling glial progenitor cells.

We investigated if these GPCs, which were both highly connected to dense collections of regulatory elements and highly enriched for lineage-defining transcription factors, might be indicators of the eventual differentiation end point, and thus possibly drive differentiation in the pseudotime trajectory. When we re-projected ATAC-seq pseudobulk samples from A, B, and C cycling population branches by only using GPC-associated chromatin signals, we observed that samples remapped to more mature expression states, which were associated with later pseudotime annotations (**Figure 6F**). In contrast, reprojections using random gene subsets or modules of genes moved non-specifically towards the center of the manifold (**Figure S15**). This observation suggests that chromatin patterns linked to GPC genes in these cycling cells already exhibit a signature of an advanced transcriptional cell state. We then projected the scRNA data from the joint multiome data set into the module-based manifold, and then transferred the corresponding scATAC cluster labels to these cells. Consistent with the singleome data, cells projecting to the cycling domain exhibited distinct accessibility signatures of more terminally differentiated cells from each branch in the same cells (**Figure 6G**). Based on these results, we propose that during corticogenesis, progenitors entering the cell cycle may be epigenetically primed toward future cell fates, and that this information is encoded specifically in GPCs, a set of genes with large numbers of linked enhancers that is enriched for lineage-defining TFs.

### Deep learning models prioritize disruptive noncoding mutations in ASD

We next aimed to use this accessibility and gene expression atlas to interpret non-coding *de novo* mutations in ASD using data from the Simons Simplex Collection, which includes a catalog of over 200,000 such mutations in 1,902 families (An et al., 2018) (**Table S20**). Naïve overlap of mutations with cluster-specific scATAC peaks produced no enrichment for mutations in ASD subject relative to those in unaffected siblings (odds ratio (OR) = 1.02 for GluN6 cluster, Fisher’s Exact Test *P* = 1.0; **Figure S16A**), indicating that peak-level annotations alone are insufficient to resolve a sparse set of causal mutations.

Deep learning models trained to relate genomic sequence to chromatin accessibility have proven useful for prioritizing disease-relevant non-coding genetic variants based on their predicted regulatory impact (Kelley et al., 2016, 2018; Zhou and Troyanskaya, 2015). We therefore trained convolutional neural networks, based on the recent BPNet architecture, to learn models that could predict base-resolution, pseudo-bulk chromatin accessibility profiles for each of our scATAC-seq derived cell types (**Figure 7A**) (Avsec et al., 2020). These models utilize DNA sequence across 1000 bp flanking each peak summit to predict 5’ Tn5 insertion counts profiles at single-nucleotide resolution (**Methods**). To correct for potential sequence composition biases, we trained the models on peak regions and genomic backgrounds matched for GC content and motif density (**Figure S16B**). The models showed high and stable correlation between total predicted and observed Tn5 insertion count coverage across all peak regions in held-out chromosomes across five-folds of cross-validated models (e.g., GluN6, mean Spearman rho = 0.58; **Figure S16C, Table S21**). Next, to predict cell context-specific effects of a candidate mutation on chromatin accessibility, we used our cluster-specific BPNet models to compute local disruption score based on the allelic fold-change in predicted counts in a 200 bp window around the mutation (**Methods**). We computed cluster-specific enrichment of high-effect size mutations in cases versus controls and observed significant enrichments (> 1.2-fold) for GluN2/3/4/6/9, as it has been previously indicated (Gandal et al., 2018; Li et al., 2018a; Parikshak et al., 2013; Trevino et al., 2020; Willsey et al., 2013). Moreover, we found an association with IN2/3/4, nIPC, late RG and early RG clusters, with early RG cluster showing the highest enrichment (OR = 1.909, excess of 20, Fisher’s exact *P* < 0.05; **Figure 7B**; **Table S22**). In contrast, BPNet models trained on human fetal heart enhancers produced no enrichment (OR = 1.01, *P* = 1.0), and naïve overlap enrichment with a set of fetal heart enhancers also produced no enrichment for case mutations (OR = 0.97, *P* = 1.0; **Figure 7C**). Together, these results illustrate the power for prioritizing putative causal non-coding mutations by utilizing mutation effect scores from base pair-resolution predictive models trained on chromatin accessibility landscapes in disease-relevant cell states.

**Figure 7:**
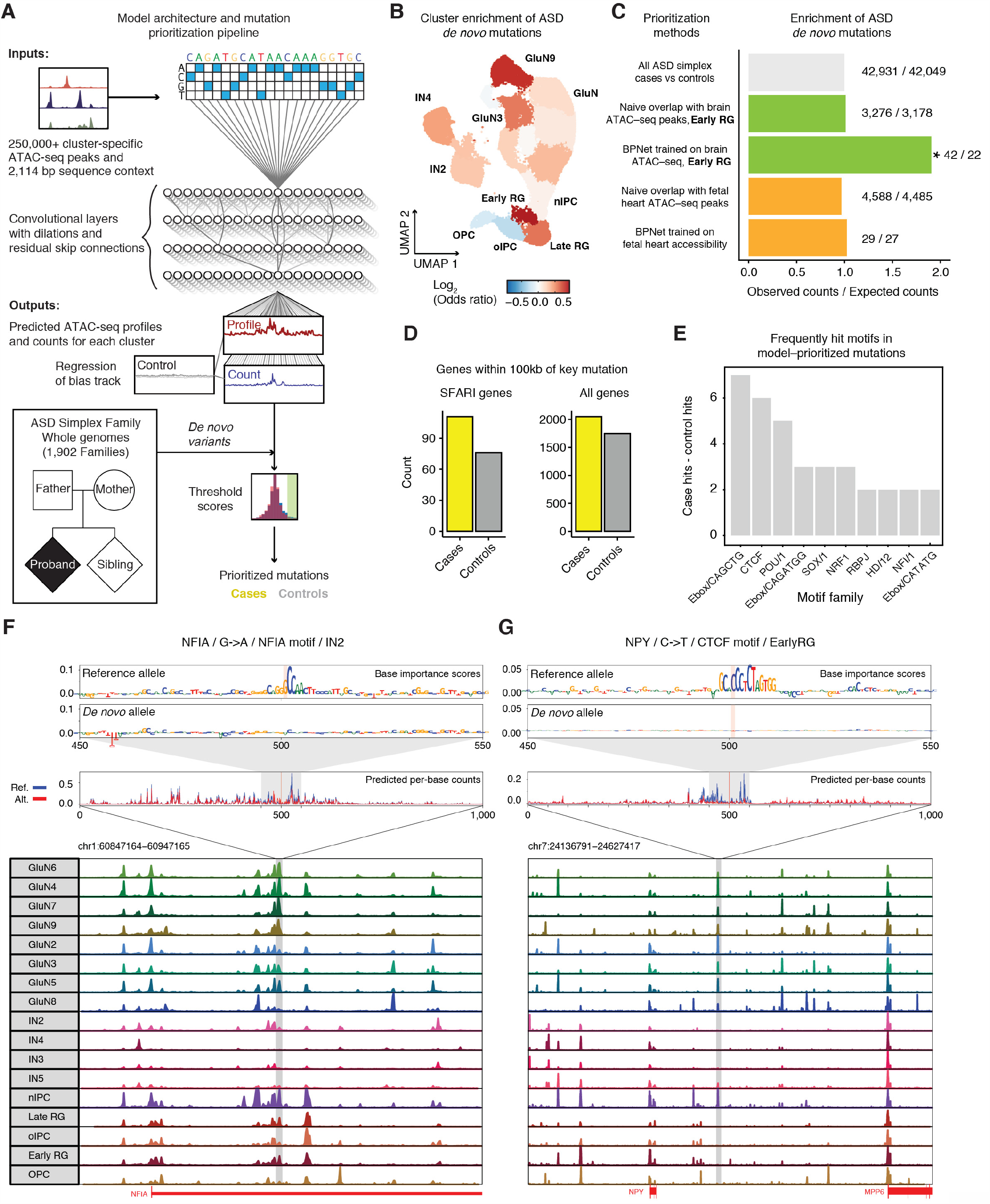
Disease association of gene regulatory elements. (A) Schematic of deep learning mutation prioritization pipeline for ASD-associated mutations from the Simons Simplex Collection (SSC). Model inputs and output, bias correction, and thresholding are shown. (B) Cluster-specific BPNet enrichments visualized in scATAC UMAP. (C) Bar plot showing the enrichment of cases versus controls using different prioritization methods. Colors represent the baseline of all cleaned SSC mutations (grey), this scATAC-seq dataset (green) and a set of fetal heart enhancers (orange). BPNet models trained on cerebral cortex scATAC enrich for case mutations; OR = 1.909, Fisher’s exact test *P* = 0.01. (D) Bar plot showing the number of prioritized mutations whose nearest gene is a SFARI gene. All SFARI genes were included. Cases (111) versus controls (76) are compared to the total number of prioritized mutations in cases (2051) versus controls (1749). (E) Bar plot showing the motifs that were most frequently disrupted in case mutations relative to control mutations. The y-axis denotes the excess of overlaps with motifs by prioritized mutations in cases and controls. (F) Example showing a disruptive case mutation in an intron of NFIA. The consensus logos show the importance of residues to predicted accessibility at the mutation in a 100 bp window flanking the mutation. Underneath, genome tracks indicate predicted per-base counts for ref (blue) and alt (red) alleles in a 1000 bp window flanking the mutation. At bottom, the *NFIA* locus is shown. Tracks display the aggregate accessibility of scATAC clusters. (G) Example showing a disruptive case mutation at the NPY locus, as above.

The case and control mutations prioritized by the BPNet models similar conservation scores and similar distances to the nearest TSS (**Figures S16D** and **S16E**), further highlighting the challenge of identifying these causal mutations by other means. Annotating the predicted high effect size mutations with their nearest genes (**Methods**), we observed a 1.4-fold enrichment for case mutations (n= 24) whose nearest gene was in the SFARI database compared to the control mutations (n = 17); **Figure 7D**). Next, we identified TF motifs that overlapped and were predicted to be disrupted by all the high-effect-size mutations from the BPNet models from all positively enriched clusters (**Methods, Figure 7E, Table S23**). We found that CTCF, which demarcates topological loop boundaries, was one of the most frequently disrupted motifs in cases versus controls. The NRF1 motif was another frequently disrupted motif. NRF regulates the GABA receptor subunit *GABRB1*, which has been previously implicated in neuropsychiatric diseases (Li et al., 2018b). Other frequently disrupted motif families in cases relative to controls included E-box/bHLH family motifs (ASCL1, NEUROD6) and homeobox family (PAX5) motifs, with more lineage-specific effects.

One highly disruptive mutation in our models was in an intron of *NFIA*, a key transcription factor active across developmental stages (**Figures 7F** and **S16F**). Loss of function mutations in this gene have previously been implicated as causal in ASD (Iossifov et al., 2014). The mutation was in a linked intronic enhancer for the *NFIA* target gene. We observed that this enhancer was specifically accessible in different types of cortical glutamatergic neuron clusters. The BPNet model for GluN6 predicts the mutation disrupting an NFIA motif, suggesting this mutation may dysregulate the NFIA gene expression via auto-regulatory feedback.

In the nIPC cluster, the BPNet model predicted a disruptive *de novo* mutation in an intergenic enhancer linked to the neuropeptide Y gene (*NPY*) whose TSS was 90 kb away from the mutation (**Figure 7G**). *NPY* is expressed in the subplate (Miller et al., 2014) and in early RG in the mid-gestation human cortex (**Figure S16G**), and genomic deletions of the NPY receptors have been associated with ASD (Ramanathan et al., 2004). The model further predicted this *de novo* mutation to disrupt a CTCF binding site at a chromatin loop anchor, suggesting a potential mechanistic impact on the chromatin architecture of this locus.

## DISCUSSION

Here, we generate paired transcriptome and epigenome atlases of corticogenesis across multiple time points during a critical period of cortical development, and we describe how molecular interactions between DNA binding factors and *cis*-regulatory elements regulate gene expression programs that ultimately drive neurogenesis and gliogenesis. Furthermore, we describe how rare non-coding, *de novo* mutations may act to disrupt this logic, linking human genetic variability to neurodevelopmental disease states.

We identified a set of genes (GPCs) whose local chromatin accessibility was predictive of expression levels using signals derived from single cells, possibly because of the large number of enhancers with accessibility that correlates with gene expression changes. These GPCs were significantly enriched for lineage-defining TFs. The large groups of enhancers linked to key cell-specific genes are evocative of other terms that have been used for similar phenomena, including “super enhancers” (Parker et al., 2013; Whyte et al., 2013) and “super-interactive promoters” (Song et al., 2020). Furthermore, chromatin accessibility of GPCs was consistent with a more differentiated cell state in a population of cycling progenitor cells. Recently, Ma et al. (Ma et al., 2020) reported a similar phenomenon by which accessibility at similarly-defined domains of regulatory chromatin delineate potential future cell states. We speculate that the coordinated regulatory effect of many enhancers of these lynchpin differentiation genes may help these lineage defining factors become more resistant to regulatory noise. We speculate that highly cooperative regulation of lineage determining trans-acting factors may be a general principle of fate determination, allowing cellular fate to be “locked in” by multiple correlated regulatory elements once the fate decision has been made. Effectively, such a cooperative transition might act as a ratchet, preventing backtracking along a differentiation landscape.

Examining the trajectories of glutamatergic neuron migration and maturation in our data, we found a molecular program that was surprisingly consistent across 8 weeks of gestation, defined by a sequence of motifs that included ASCL1, GLI3, SOX family, EOMES, NFI family, POU3F3, NEUROD2 and MEF2C. Differences in neuronal regulatory activity across pseudotime were more pronounced than differences between developmental stages. We further found distinct patterns of co-accessibility and regulatory interactions between TFs early in pseudotime, whereas gene TFs appeared to act more independently later in pseudotime.

Moreover, we found substantial sharing of TF-regulated gene expression programs amongst glial cells by decomposing these programs into overlapping modules. Notably, we found substantial overlap between gene modules containing canonical markers for astrocytes and oligodendrocytes, suggesting a lineage relationship. We validated the co-expression of several of these genes *in situ* in human cerebral cortex. A similar relationship was true for modules associated with astrocytes and ependymal cells. We also provided evidence for the existence of two lineages of astrocyte-like glial precursors at or before PCW24, which could align with diversity of these cells in primates (Vasile et al., 2017). Although glial modules were broadly interconnected, we found that the chromatin activity of GPCs in cycling cells was predictive of specific differentiated states, suggesting that progenitors entering the cell cycle are already primed towards distinct lineages.

Finally, our map of chromatin regulation across these distinct cell types provided a rich data set for training interpretable, cell-type specific deep-learning models that link DNA sequence to chromatin accessibility. These models can be used to read the potential regulatory impacts of de novo mutations, allowing the prioritization of high impact noncoding mutations and generating strong enrichments of mutation occurrence in cases over controls. The modeling of the regulatory potential of individual base pairs at different stages of development was crucial to enable the identification of these putative causal mutations, as simple overlap with open chromatin regions did not provide the required specificity. Indeed, using our model, we observed enrichments of mutations in cases versus controls that approached levels observed for deleterious protein-coding mutations (An et al., 2018). Furthermore, combining these models with our single-cell atlas allows for the principled interpretation of where in development highly disruptive mutations tend to occur. We anticipate that as more large-scale ATAC-seq and RNA-seq data sets across development become available, similar approaches will provide the means to accurately interpret the gene-regulatory impact of non-coding de-novo mutations associated with a broad diversity of other developmental disorders.

## ACKNOWLEDGEMENTS

We thank members of the Greenleaf, Paşca, Kundaje, and Chang labs for discussion and advice, especially B. Parks, F. Birey, J. Granja and A. Banerjee.

## AUTHOR CONTRIBUTIONS

A.E.T., F.M., J.A., S.P.P, and W.J.G. conceived the project and designed experiments. A.E.T. and F.M. performed data analysis. J.A. guided the biological interpretation of the analysis and performed validations. L.S. trained deep learning models and performed the analysis on disease relevance with assistance from A.E.T., A.S., K.F., A.Ku. and W.J.G. A.M.P. and J.A. processed the samples for single cell experiments. A.E.T. and A.Ka. performed single cell experiments. A.E.T., F.M., J.A., L.S., S.P.P. and W.J.G. wrote the manuscript with input from all authors. S.P.P and W.J.G supervised the work.

## MATERIALS AND METHODS

### Human tissue

Human brain tissue was obtained under a protocol approved by the Research Compliance Office at Stanford University. Cortical tissue was processed immediately after arrival.

### Single cell dissociation and single cell RNA-seq data generation

Dissociation of human tissue into single cells was performed as described with some modifications (Sloan et al., 2017; Trevino et al., 2020). Briefly, tissue was chopped and incubated in 30 U/ml papain enzyme solution (Worthington, LS03126) and 0.4% DNase (12,500 units/ml; Worthington, LS002007) at 37 °C for 45 minutes. After digestion, samples were washed with a protease inhibitor solution and gently triturated to achieve a single cell suspension. Cells were resuspended in 0.02% BSA/PBS and passed through a 70 μm filter before proceeding to single-cell sample preparation. Single-cell libraries were prepared using the RNA 3’ v3 protocol (10x Genomics), loading 7,000 cells per lane.

### ATAC-seq data generation

For ATAC-seq, nuclei were prepared on ice or in a centrifuge at 4 °C. All centrifugation steps were run for 5 minutes at 500 x g. 100,000 dissociated cells were washed in ice-cold ATAC-seq resuspension buffer (RSB, 10 mM Tris pH 7.4, 10 mM NaCl, 3 mM MgCl_2_), spun down, and resuspended in 100 µL ATAC-seq lysis buffer (RSB plus 0.1% NP-40 and 0.1% Tween-20 (Thermo Fisher). Lysis was allowed to proceed on ice for 5 minutes, then 900 µL RSB was added before spinning down again and resuspending in 50 µL 1X Nuclei Resuspension Buffer (10x Genomics). A sample of the nuclei was stained with Trypan Blue and inspected to confirm complete lysis. If necessary, cell concentrations were adjusted prior to starting single-cell droplet generation with the ATAC-seq NextGEM kit (10x Genomics). 4,000 nuclei were loaded per lane.

### Multiome data generation

For multiome single cell data, nuclei were prepared as above for ATAC-seq with minor changes. Specifically, 0.01% digitonin was added to the lysis buffer, and 2 U/µL RNAse inhibitor (Roche) was added to all nuclei preparation buffers. After nuclei preparation, droplets and single cell libraries were prepared using the Single Cell Multiome ATAC + Gene Expression kit (10x Genomics) and 4,000 nuclei were loaded per lane.

### scRNA processing

Raw sequencing data were converted to fastq format using the command ‘cellranger mkfastq’ (10x Genomics, v.3.1.0). scRNA-seq reads were aligned to the GRCh38 (hg38) reference genome and quantified using ‘cellranger count’ (10x Genomics, v.3.1.0). ‘Velocyto’ (v.0.17.17) (La Manno et al., 2018) was used to obtain splicing-specific count data for downstream RNA velocity analysis.

Count data was further processed using the ‘Seurat’ R package (v.3.1.4) (Stuart et al., 2019), using Gencode v.27 for gene identification. We removed cells with less than 500 informative genes expressed, cells with less than 500 sequenced fragments and cells with more than 40% of counts corresponding to mitochondrial genes. Genes not contained in the Gencode annotation were excluded from further analysis. We performed doublet analysis using the ‘DoubletFinder’ R package (v.2.0.2) (McGinnis et al., 2019), but did not find clear evidence of cell doublets biasing our unsupervised analysis and therefore did not apply doublet filtering. Count data was log-normalized and scaled to 10,000. PCA analysis was based on the 2,000 most variable genes. The top 50 principal components (PCs) were retained for further analysis, excluding one component because it was strongly associated with the expression of more than 5 genes related to cell stress (*HSPA, JUN, FOS, DUSP* gene families). Nearest neighbors were computed based on the PC representation, and 23 clusters were identified using Louvain clustering implemented in Seurat’s ‘FindClusters’ function (‘resolution=0.5’). 2-dimensional representations were generated using uniform manifold approximation and projection (UMAP) (McInnes et al., 2020) as implemented in Seurat and the ‘uwot’ R packages (v.0.1.8; parameter settings: ‘min.dist=0.8’, ‘n.neighbors=50’, ‘cosine’ distance metric).

### scATAC processing

Raw sequencing data were converted to fastq format using ‘cellranger-atac mkfastq’ (10x Genomics, v.1.2.0). scATAC-seq reads were aligned to the GRCh38 (hg38) reference genome and quantified using ‘cellranger-atac count’ (10x Genomics, v.1.2.0).

Fragment data was further processed using the ‘ChrAccR’ R package (v.dev.0.9.11+). We filtered out cells with less than 1,000 or more than 50,000 sequencing fragments. TSS enrichment was computed as a metric of signal-to-noise ratio using methods described in (Granja et al., 2019) and we discarded cells with a TSS enrichment less than 4. Fragments on sex chromosomes and mitochondrial DNA were excluded from downstream analysis.

In order to obtain a low dimensional representation of single-cell ATAC datasets in terms of principal components and UMAP coordinates, we applied an iterative latent semantic indexing approach (Granja et al., 2019). This approach also identified 22 cell clusters and a consensus set of 657,930 cluster peaks. In brief, in an initial iteration clusters were identified based on the 20,000 most accessible 5kb-tiling regions. Here, the counts were first normalized using the term frequency - inverse document frequency (TF-IDF) transformation (Cusanovich et al., 2018), and singular values were computed based on these normalized counts. Initial clusters were identified based on the top 25 singular values using Louvain clustering (as implemented in the Seurat package, resolution parameter = 0.6), excluding the first singular value as it exceeded a correlation coefficient of 0.5 with read depth. Peak calling was then performed on the aggregated insertion sites from all cells of each cluster using MACS2 (v2.1.1). A consensus set of peaks uniform-length non-overlapping peaks was obtained by selecting the peak with highest score from each set of overlapping peaks. In a second iteration, the 50,000 peaks whose TF-IDF-normalized counts exhibited the highest variability across the initial clusters provide the basis for a refined clustering using the top 50 derived singular values. In the final iteration, the 50,000 most variable peaks across the refined clusters were identified as the final peak set and singular values were computed again. UMAP coordinates and ATAC clusters were determined based on the top 10 of these final singular values. 2-dimensional representations were generated using UMAP as implemented in the ‘uwot’ R package (v.0.1.8; parameter settings: ‘min.dist=0.6’, ‘n.neighbors=50’, ‘cosine’ distance metric).

ChromVAR (Schep et al., 2017) (v.1.6) was used to obtain TF accessibility profiles using position weight matrices from the JASPAR 2018 database (Khan et al., 2018). Gene activity scores were computed as the aggregated accessibility of TSS-associated peaks using ‘ChrAccR’. For this, counts in peaks within 100,000 bp of a TSS have been summed up using weights assigned by a radial basis function (RBF) with a width parameter sigma=10,000 bp, setting a minimum asymptotic weight of 0.25. For each gene, the resulting scores were normalized by the sum of the weights. For visualization and downstream analysis counts from single-cells have been rescaled to 10,000 counts and have been log_2_-normalized. For enhanced visualization in 2-dimensional UMAP space, gene activity scores have been smoothed using the MAGIC diffusion algorithm (van Dijk et al., 2018) with cell neighborhoods determined in singular value space.

We created ATAC signal tracks by summing insertion counts in cluster pseudobulk samples in 200bp genomic tiling windows and provide trackhub compatible with the WashU Epigenome Browser (http://epigenomegateway.wustl.edu) containing these profiles in addition to inferred CRE-gene links.

### Multiome data processing

Raw sequencing data were converted to fastq format using ‘cellranger-arc mkfastq’ (10x Genomics, v.1.0.0). scATAC-seq reads were aligned to the GRCh38 (hg38) reference genome and quantified using ‘cellranger -arc count’ (10x Genomics, v.1.0.0).

RNA count data was further processed using ‘Seurat’ as described above, with the exception that all 50 principal components were retained. This resulted in 9,818 cells after filtering, which were assigned to 14 clusters in the unsupervised analysis. ATAC fragment data was further processed using ‘ChrAccR’ as described above, resulting in 9,091 cells post-filtering, assigned to 16 clusters and a consensus peak set of 467,315 elements. Jointly applying ATAC and RNA filters resulted in 8,981 cells with high-quality measurements across both modalities.

Processed data and additional analysis code can be found at https://github.com/alexandrotrevino/brainchromatin.

### Matching of single-cell transcriptomes and epigenomes

Canonical correlation analysis (CCA) as implemented in Seurat has been applied to matched single-cell RNA and ATAC data from each gestational time point individually. For this purpose, we computed log-normalized and scaled gene activity scores as surrogates for gene expression in the cells profiled by scATAC-seq. As integration features, we used the union of the 2,000 most variable genes in each modality as input to Seurat’s ‘FindTransferAnchors’ function with reduction method ‘cca’ and parameter ‘k.anchor=10’. For each cell profiled by scRNA-seq and each cell profiled by scATAC-seq we identified the nearest neighbor cell in the respective other modality by applying nearest-neighbor search in the joint CCA L2 space. Nearest neighbors were determined using the ‘FNN’ R package employing the ‘kd_tree’ algorithm with Euclidean distance. These nearest-neighbor-based cell matches from all gestational time points were concatenated to obtain dataset-wide cell matches across both modalities.

### Linking gene regulatory elements and gene expression across all cell types

We identified peak-to-gene links using a correlation-based approach (Corces et al., 2018) applied to pseudobulk samples aggregating scATAC and scRNA counts. These pseudobulk samples were defined by randomly sampling 200 cells from the entire scATAC-seq dataset. These 200 seed cells were combined with their respective 99 nearest neighbor cells in ATAC-PC space, such that each pseudobulk sample comprised 100 cells in total. Pseudobulk ATAC insertion counts for peaks were obtained by summing peak insertion counts across the respective single-cell members. Matching RNA cells were obtained by selecting the 100 scRNA cells that resembled nearest neighbors to the 100 ATAC cells in CCA space. Pseudobulk RNA gene counts were obtained by summing gene counts across the respective single-cell members. Similarly in the multiome dataset, 200 pseudobulk samples of 100 cells each were sampled from the ATAC modalitity, and the same cells were aggregated in RNA space. Each matched pseudobulk sample was annotated with the majority cluster and age assignments of its contingent RNA and ATAC cells respectively.

We then obtained candidate peak-gene pairs by associating peaks with a genomic distance between 1 and 250 kb to the TSS of protein coding and lincRNA genes to the respective genes. For each candidate peak-gene pair we computed the Pearson correlation coefficient of CPM-normalized counts of accessibility and gene expression data and computed FDR-adjusted *P*-values for these coefficients based on their *t*-statistic. We defined a set of 64,878 high-confidence peak-to-gene links by only retaining pairs with |PCC| > 0.4 and FDR-adjusted *P*-value < 0.05. Using the same method, a corresponding set of 76,374 links was obtained for the multiome data. Overlap between inferred and multiome peak-gene links was computed by creating “GenomicInteraction” objects for each, with the peak as the first anchor and the gene promoter as the second, then applying the function ‘findOverlaps’ with parameter “use.region = ‘both’”.

### Projection of external datasets into the scRNA landscape

We retrieved scRNA data from the developing human cerebral cortex (Bhaduri et al., 2020). We downloaded the normalized data from the UCSC Cell Browser (https://cells.ucsc.edu; dataset ID: ‘organoidreportcard/primary10X’) and the data was read into a Seurat object using custom R scripts. We then projected the data into our scRNA UMAP space using the ‘uwot’ model stored in our dataset, i.e. we used an identical principal component gene loadings and ‘uwot’ model parametrization. This UMAP space representation allowed us to assign a nearest neighbor from our scRNA cells to each cell in the external dataset. Cell annotation (pseudotime, cell cluster, etc.) were transferred from these nearest neighbors. Jaccard indices were computed between the transferred annotation and the downloaded external metadata.

Similarly, we downloaded 10x Genomics scRNA data from the Allen Brain Map (https://portal.brain-map.org/atlases-and-data/rnaseq). The downloaded raw count data was read into a Seurat object and processed using the same steps and parameters used for processing our scRNA data. Projection and annotation transfer were done in the same way as for the external developing brain dataset. For Figure S7, we restricted the projection to cells labelled as excitatory neurons (‘Exc’) in the external cell metadata.

### Projection of multiome data into the scRNA and scATAC landscapes

Based on the RNA-based gene counts, we projected the multiome data into our scRNA UMAP space using the ‘uwot’ model stored in our scRNA dataset, i.e. we used an identical principal component gene loadings and ‘uwot’ model parametrization. Similarly, multiome cells were projected into scATAC UMAP space based on the ‘uwot’ model derived from the scATAC dataset using the same peak loadings. We used these projections to assign a nearest neighbor from our scRNA cells or scATAC cells to each cell in the multiome dataset. Cell annotation (pseudotime, cell cluster, etc.) were transferred from these nearest neighbors.

### Identification of genes with predictive chromatin (GPCs)

The definition of GPCs is primarily based on high gene activity-expression correlations across single cells. To make this analysis more robust to technical variation, we restricted our analysis to the most variable genes across dorsal forebrain cells (1999 genes). Specifically, we used the “findVariableGenes” function from the URD package with parameters “diffCV.cutoff = .15, mean.min = 0.004” (Farrell et al., 2018). For each variable gene, we computed Spearman’s correlation coefficients between the vector of gene activity scores for ATAC cells and the vector of expression scores in the corresponding nearest neighbor cells in RNA data. We also compared these correlations to the number of linked enhancers per gene (see above). From this subset, we defined GPCs as genes in the top 10% of gene activity-expression correlations that were linked to a minimum of 10 CREs.

### Definition of RNA velocity and pseudotime in excitatory neuron trajectories

Excitatory neuron trajectories were defined based on RNA cells in selected clusters (cf. Table S6). We computed RNA velocity using custom R scripts interfacing with the ‘scVelo’ toolkit (v.0.1.25) (Bergen et al., 2020) via the ‘reticulate’ R-Python interface. For this, we exported the Velocyto-derived spliced and unspliced counts along with Seurat-derived PC and UMAP representations of single cells as ‘AnnData’ objects. We filtered the dataset using the scVelo function ‘pp.filter_and_normalize’ (parameters: min_shared_counts=10, n_top_genes=2,000) and computed moments using ‘pp.moments’ (n_pcs=30, n_neighbors=30). We then used ‘tl.velocity’ with mode=‘stochastic’ to compute cell velocities and ‘tt.velocity_graph’ to compute a velocity graph. Potential root and end point cells for the trajectory were computed using ‘tt.terminal_states’. To compute cell pseudotime scores, we employed a modified version of the scVelo function ‘tt.velocity_pseudotime’. In contrast to the original version of the function which combines diffusion estimates from a forward pass starting in the root cells and a backwards pass starting in the end point cells, this modified version only applies the forward pass starting in the root cells. This was necessary because scVelo-identified end point cells that were inconsistent with our notion of trajectory. We re-imported the scVelo-derived cell annotations (velocity vectors, pseudotime, root and end point probabilities) into the metadata of the R-based Seurat objects. Finally, cell pseudotime scores were rescaled to their quantiles using the R function ‘ecdf’.

Additionally, in order to quantify when in pseudotime a gene is expressed we computed a weighted average pseudotime value. We define this ‘gene pseudotime’ for each gene *j* as

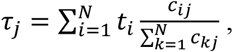

where *N=363* is the number pseudobulk samples used for linking regulatory elements to genes (see below), *t*_*i*_ is the mean pseudotime across all cells in pseudobulk sample *i*, and *c*_*ij*_ is the aggregate RNA count for pseudobulk sample *i* in gene *j*.

Pseudotime of cells profiled using scATAC-seq were defined as the pseudotime of their nearest RNA-cell neighbor in CCA space.

### Linking gene regulatory elements and gene expression in the excitatory neuron trajectory

To facilitate aggregate analysis along pseudotime, we obtained pseudobulk samples by sorting cells based on their pseudotime scores and merging bins of 100 cells. The same correlation-based approach as used on all cell types (see above) was applied to these pseudobulk samples, linking peaks to cluster-specific genes. These cluster-specific genes were identified from the scRNA data of cells included in the excitatory neuron trajectory employing a Wilcoxon test as implemented in Seurat’s ‘FindAllMarkers’ function and applying thresholds of 0.01 and 0 for test-derived *P*-values and log(fold-changes) respectively. We retained links with accessibility-gene expression correlation coefficients with PCC > 0.4 and FDR-adjusted *P*-value < 0.05, which resulted in 13,989 high-confidence positively correlated peak-to-gene links with specificity to the excitatory neuron trajectory. These links were clusters using k-means (k=5) clustering based on the z-score-scaled expression levels of the associated genes. Enrichment analysis for these clusters were performed using the ‘topGO’ (v.2.36.0) R/Bioconductor package (Gene Ontology enrichment), Fisher’s exact tests on manually curated gene sets and Fisher’s exact tests as implemented in the R/Bioconductor package ‘LOLA’ (Sheffield and Bock, 2016) (v.1.14.0) for peak TF motif occurrences (based on genome-wide scans of JASPAR 2018 PWMs using the ‘motifmatchr’ R package).

### Matching TF motifs to expressed TF genes in the excitatory neuron trajectory

To avoid correlation biases in closely-related TF motifs, we used a database of previously annotated clusters of putative binding motifs (Vierstra et al., 2020). For each motif cluster, we computed the pairwise Pearson correlation coefficients between chromVAR motif activity scores (computed from the annotated genome-wide sites of that cluster) and gene expression of all genes attributed to motifs in that cluster (Figure 3H). These PCCs were computed across the same pseudotime-pseudobulk samples that were used for CRE-gene linking. We then matched each gene to the motif cluster that exhibited the highest correlation with that gene (Figure 3F). We identified 24 dynamic motifs clusters representing 31 TFs whose gene loci are linked with at least one CRE and that exhibit high correlation coefficients of motif cluster activity and TF expression (PCC ≥ 0.4) for downstream analysis (Figure 3F–J).

### Calculation of motif synergy and correlation scores

We used chromVAR to compute synergy and correlation scores for the above 24 dynamic motifs clusters. (Schep et al., 2017). We used the ‘getAnnotationSynergy’ chromVAR function to compute synergy scores, which represent the excess variability of chromatin accessibility in CREs that contain binding sites from two different motif clusters compared to a random sub-sample of CREs which contains binding sites from only one of the motif clusters (the one with greater variability). It thus suggests a co-dependence of TFs belonging to the two motif clusters. In order to assist in the discrimination between this co-dependence and co-expression, we also computed motif correlation coefficients using the ‘getAnnotationCorrelation’ function in chromVAR, defined as the correlation between the deviation scores for the CREs that only contain binding sites from only one or the other motif cluster.

### Inference of TF regulatory networks

We established a network of TF regulatory linkages by testing whether CREs with TF motif occurrences exhibited significantly better peak-to-gene linkages than CREs without the motif. In this network the nodes correspond to the 31 dynamic TFs in the excitatory neuron trajectory. We draw a directed edge between TF1 and TF2 iff the regulatory elements linked to TF2 that contain binding sites for the motif cluster that TF1 belongs to exhibit significantly larger correlation coefficients than regulatory elements that do not contain a binding site for TF1 (one-side Wilcoxon Rank Sum test *P*-value < 0.01).

### Fuzzy c-means: clustering and re-projection approach

For fuzzy clustering analysis, 1,267 seed cells were first selected at random from glial clusters (10% of single cells), with the number selected proportional to the cluster size. Pseudobulk data sets were sampled by combining these cells with their 50 nearest neighbors in scRNA PCA space. Next, 1,957 variably expressed genes were determined using the function ‘findVariableGenes’ from the R package ‘URD’. A pseudobulk counts matrix was made by summing feature counts across the respective single cell members comprising each aggregate.

Fuzzy c-means clustering was performed on this pseudobulk matrix using the function “cmeans” from the R package ‘e1071’ with parameters c = 14 and m = 1.25, resulting in a gene-by-module “membership matrix” and a sample-by-module “centers matrix”. To determine a ‘fixed’ or binarized module membership for downstream analyses, we defined a threshold membership score as the maximum score at which all genes were assigned to a cluster (threshold = 0.06). Gene ontology enrichments for each module were computed using the function ‘enrichGO’ from the R package ‘clusterProfiler’. Module connectivity was computed between all module pairs using the Jaccard index, and modules were linked by applying a threshold of 0.2 of the Jaccard index of gene sharing. This threshold was chosen by applying the elbow method. To visualize the connections between modules, the centers matrix (sample-by-module) was used as the basis for dimensionality reduction with UMAP, using the R package ‘umap’.

Finally, this process was repeated, sweeping the clustering parameters (c, m) and the membership threshold across a range of values; from c = 6 to c = 30, and from m = 1 to 2; to ensure that the structure of the resulting embedding was not overly sensitive to the clustering parameters.

### Projecting ATAC-seq data into fuzzy clustering space

Pseudobulk samples of scATAC cells were generated using the same approach described above for gene activity scores. This matrix was subsetted to match the features (genes) of the RNA fuzzy clustering analysis. In the case of missing features, values were imputed using their median gene activity. To project ATAC-seq cells into the RNA fuzzy clustering embedding, we transposed the membership matrix and multiplied it with the gene activity-by-pseudobulk matrix. Finally, we used the “predict” function in R ‘stats’, with the fuzzy clustering UMAP model as the first argument, and the resulting transposed product matrix as the second, to determine the UMAP coordinates of ATAC pseudobulks.

### Differential branch activity analysis

Branches were defined by grouping ATAC-seq pseudobulks projecting into the early part of the fuzzy clustering UMAP (into the Cyc cluster) according to their full-dataset cluster annotation, resulting in three branches. Differential gene activities were calculated using Wilcoxon rank sum tests to compare branch A to B and C, B to A and C, and C to A and B. Genes for each branch were ranked by their average log_2_ fold change in the differential test. The 50 most unique genes for each branch were visualized in a row-scaled heatmap.

Gene set enrichment analysis of GPCs was performed using the Kolmogorov-Smirnov test for GPC ranks in the differential test, relative to non-GPC ranks.

Motif enrichments for GPC TFs were derived by computing a Chi-square test for the enrichment of motifs in peaks linked to differential gene activities. To find the TF motifs that best correspond to GPC TF genes, the best-correlated TF motif activity (chromVAR) for each GPC TF across glial pseudobulks was used.

### Characterization of astrocyte heterogeneity

We computed motif enrichments between peaks linked to modules 13 and module 14, which both contained *AQP4, APOE*, and *ALDH1* as members, using a chi-squared test. Resulting *P*-values were adjusted for multiple testing using a Bonferroni correction. Next, to define groups of astrocyte cells (samples in contrast to astrocytic gene signatures (modules)), we re-clustered the RNA-derived glial pseudobulk samples, and performed unbiased differential expression testing using “DESeq2” between clusters c0 and c5, which highly expressed astrocyte genes (Zhang et al., 2016). A stringent FDR (Benjamini-Hochberg) of 1e-20 was invoked to call differential genes, since applying the DESeq2 (Love et al., 2014) framework to pseudobulks deflated *P-*values. The top 200 most differential genes were used to plot aggregate differential gene expression within the alternative dataset.

### *De novo* Mutation Filtering

*De novo* mutations from 1902 individuals with ASD and their unaffected siblings from the Simons Simplex Collection were obtained from (An et al., 2018). Mutations of coding or splice consequence, as annotated by GENCODE v27 (https://www.gencodegenes.org/human/release_27.html), were ignored from final analysis. Additionally, *de novo* mutation calls that are observed in gnomAD (Karczewski et al., 2020), in nonstandard chromosomes, within the low complexity repeat regions from the UCSC browser table RepeatMasker (http://hgdownload.cse.ucsc.edu/goldenpath/hg38/database/rmsk.txt.gz), were removed from downstream analysis. Also, de novo mutations appearing in both affected and unaffected siblings and multiple SSC families (i.e., non-singleton *de novo* mutations) were removed.

### BPNet deep learning models to predict base-resolution cluster-specific pseudo-bulk scATAC-seq profiles from DNA sequence

BPNet is a sequence-to-profile convolutional neural network that uses one-hot-encoded DNA sequence (A=[1,0,0,0], C=[0,1,0,0], G=[0,0,1,0], T=[0,0,0,1]) as input to predict single nucleotide-resolution read count profiles from assays of regulatory activity (Avsec et al., 2020). The models take in a sequence context of 2,114 bp around the summit of each ATAC-seq peak, and predicts cluster-specific scATAC-seq pseudobulk Tn5 insertion counts at each base pair for the central 1,000 bp. The BPNet model also uses an input Tn5 bias track which is concatenated to the prefinal layer as explained below. Our BPNet model is based on the architecture introduced in (Avsec et al., 2020). The model architecture consists of 8 dilated residual convolution layers, with 500 filters in each layer. At each layer, the Keras Cropping 1D layer is used to clip out the two edges of the sequence, to match the inputs concatenated to the output of each convolution, which naturally trims the 2,114 bp sequence to a final 1,000 bp profile. Each dilated convolutional layer has a kernel width of 21 and the dilation rate is doubled for every convolutional layer starting at 1. The model predicts the base-resolution 1,000 bp length Tn5 insertion count profile using two complementary outputs: (1) the total Tn5 insertion counts over the 1,000 bp region, and (2) a multinomial probability of Tn5 insertion counts at each position in the 1,000 bp sequence. The predicted (expected) count at a specific position is a multiplication of the predicted total counts and the multinomial probability at that position. To predict the total counts in the 1,000 bp window, the output from the last dilated convolutional layer is passed through a GlobalAveragePooling1D layers in Keras. We estimate the “tn5 bias” for the input sequence using the TOBIAS method (Bentsen et al., 2020). This total bias is concatenated with the output of the pooling layer and passed through a Dense layer with 1neuron to predict total counts. To predict the per-base logits of the multinomial probability profile output, the output from the last dilated residual convolution is appended with per base TOBIAS “tn5 bias” and passed through a final convolution layer with a single kernel and a kernel width of 1 to predict the per-base logits. BPNet uses a composite loss function consisting of a linear combination of a mean squared error (MSE) loss on the log of the total counts and a multinomial negative log-likelihood loss (MNLL) for the profile probability output. We use a weight of [16.1, 12.5, 10.8, 12.4, 13.3, 7.4, 11.6, 13.1, 9, 7.5, 4.7, 7.2, 4.9, 16.1, 4.2 & 2.3] for the MSE loss for clusters c0–c15, and a weight of 1 for the MNLL loss in the linear combination. The MSE loss weight is derived as the median of total counts across all peak region for each cluster divided by a factor of 10 (Avsec et al., 2020). We used the ADAM optimizer with early stopping patience of 3 epochs.

A separate BPNet model was trained on pseudobulk scATAC-seq profiles from each scATAC-seq cluster with > 500 cells (to ensure sufficient coverage for reliable training). We used a 5-fold chromosome hold-out cross-validation framework for training, tuning and test set performance evaluation. The training, evaluation and test chromosomes used for each fold are as follows.

Test chromosomes: fold 0: [chr1], fold 1: [chr19, chr2], fold 2: [chr3, chr20], fold 3: [chr13, chr6, chr22] & fold 4: [chr5, chr16] Validation chromosomes: fold 0: [chr10, chr8], fold 1: [chr1], fold 2: [chr19, chr2], fold 3: [chr3, chr20] & fold 4: [chr13, chr6, chr22].

The model’s performance was evaluated using two different metrics for the two output tasks separately. For the total counts predicted for the 1,000 bp region, the model’s performance is computed with the Spearman correlation of predicted counts to actual counts. The per-base read count track is evaluated using the Jensen-Shannon divergence distance, which computes the divergence between two probability distributions; in this case the actual per base read profile for the 1,000 bp region and the predicted per base read profile for the 1,000 bp region.

We initially trained the BPNet models only on cluster-specific scATAC-seq peaks, without including other background regions. Scoring the ASD *de-novo* mutations (see below) using these models trained only on positive peaks, we observed a significant shift in model disruption scores for the case mutation when compared to the control mutations (Cluster C10 fold 0 model rank-sum *P*-value: 6.2e-07). Given that the odds ratio of loss of function (LoF) mutations between cases and control in ASD is 2.09 (n-cases= 81, n-controls= 38, Fisher’s exact *P*-value= 0.0001) and majority of the *de novo* mutations are autosomal dominant, we expect a very small fraction of the mutations to be disrupting regulation. We observed a slight difference in G/C composition within 2 kb around the case mutations compared to control mutations (Ranksum *P*-value: 0.02826). To avoid any potential biases or confounding due to G/C content in the sequence, for every cluster specific peak sequence, we randomly sampled background regions from the genome with matched G/C content and regions that had similar motif density called using JASPAR motif scan. We retrained the cluster specific BPNet models on a larger set of regions that included the peaks and 10% of matched background regions. The new models that included G/C matched background regions in the training set did not exhibit a systematic shift in the mutation disruption score distribution between cases and controls (Cluster C10 fold0 G/C matched background trained Ranksums *P*-value: 0.273). Hence, we used these models for all the subsequent analyses. Code for training BPNet models as well as the saved models are available at https://github.com/GreenleafLab/Brain_ASD

### Scoring impact of ASD *de novo* mutations on cluster-specific chromatin accessibility using neural network models

The filtered *de novo* mutations from affected (cases) and unaffected siblings (controls) described in the previous section, were first overlapped with the peak regions from each scATAC-seq cluster. We then used the cluster-specific BPNet models to predict the allelic impact of all mutations overlapping cluster-specific peak regions. For each mutation, we used the BPNet model to predict the base-resolution read count profile corresponding to the input sequence (2,114 bp) containing the reference allele of the mutation at its center. We then used the model to predict the base-resolution read count profile corresponding to the input sequence (2,114 bp) containing the alternate allele of the mutation at its center. Using these two profiles, we compute the impact score of the mutation as the sum of the allelic difference in the per-base read counts over a 100 bp window around the mutation using the formula:

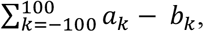

Where

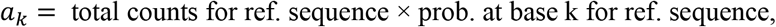

And

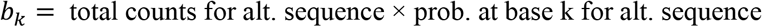

For each mutation, the impact scores were computed and averaged over cluster specific BPNet models trained on each of 5 folds. We prioritized high impact mutations as those that have an impact score > 20 in any scATAC-seq cluster.

### Cluster-specific enrichments of mutations in ASD cases versus controls

We computed a 2 x 2 contingency table to estimate overlap enrichments of case versus control mutations with cluster-specific peaks. The first axis splits all *de novo* mutations based on whether they were found in cases versus controls. The second axis splits all *de novo* mutations based on whether they overlap a cluster-specific peak. The enrichment *P*-value and odds-ratio were computed using Fishers Exact Test available in the SciPy package in Python.

We used a similar procedure to estimate enrichment of case versus control *de novo* mutations with BPNet’s classification of mutations as high impact or low impact. In this case, the first axis of the 2 x 2 contingency table splits all *de novo* mutations based on whether they were found in cases versus controls. The second axis splits all *de novo* mutations based on whether they are predicted to have a high impact score (> 20) or not using a cluster specific BPNet. High impact score mutations are pre-filtered to always overlap cluster-specific peak regions.

### Enrichment analysis of transcription factor motifs predicted to be disrupted by high impact mutations

We identified predictive nucleotides in the sequence of each peak region in each scATAC-seq cluster through the lens of the corresponding cluster-specific BPNet models using DeepSHAP (https://shap.readthedocs.io/en/latest/generated/shap.DeepExplainer.html) a specific implementation of the DeepLIFT method (Shrikumar et al., 2019). DeepLIFT decomposes the predicted output of a BPNet model for a specific input sequence into contribution (importance) scores of each base in the input sequence relative to a reference input of dinucleotide shuffled sequences. We compute per-base DeepLIFT scores for the sequences of all peaks in each scATAC-seq cluster using the total count output of the cluster-specific BPNet model. We computed separate DeepLIFT scores for sequences with reference and alternate alleles of mutations. Nucleotides with large changes in DeepLIFT scores for the two sequences represent subsequences that are significantly impacted by the mutation. These subsequences typically overlap transcription factor motif instances.

We overlapped all high impact case and control mutations with genomic instances of transcription factor motifs from the previously annotated database (Vierstra et al., 2020). Given that multiple motifs can overlap a mutation, we resolved these cases by leveraging DeepLIFT scores from BPNet models corresponding to the scATAC-seq cluster for which the mutation had the highest impact score. We computed the mean per-base DeepLIFT importance scores over each motif instance overlapping the prioritized mutations. The motif with the highest score was picked as the motif hit for the prioritized mutation for every motif overlap observed.

### Immunohistochemistry

Immunohistochemistry was performed as described (Trevino et al., 2020). Briefly, PCW17 and PCW21 human cortical tissue was fixed overnight at 4°C in 4% paraformaldehyde (PFA, Electron Microscopy Sciences). Samples were then washed with PBS and transferred to a 30% sucrose solution for 48-72 hours, then embedded in OCT (Tissue-Tek OCT Compound, 4583, Sakura Fenetek) and 30% sucrose at a 1:1 ration, and snap-frozen in dry ice. Cryosections were obtained using a cryostat (Leica) set at 30 μm and mounted on Superfrost Plus Micro slides (VWR, 48311-703). Next, sections were blocked and permeabilized for 1 hour at room temperature in blocking solution (10% normal donkey serum, 0.3% Triton-X in PBS) and incubated with primary antibodies diluted in the same solution overnight at 4°C. The following primary antibodies were used: anti-ASCL1(Mouse, 1:100, BD Biosciences, 556604), anti-CTIP2 (Rat, 1:300, Abcam, ab18465), anti-EGFR (Rat, 1:200, Abcam, ab231), anti-GFAP (Rabbit, 1:1,000, Dako, Z0334), anti-GFAP (Rat, 1:1000, Thermo Fisher Scientific, 13-0300), anti-HOPX (Mouse, 1/50, Santa Cruz, sc-398703), anti-KI67 (Mouse, 1:500, BD Biosciences, 550609), anti-OLIG2 (Rabbit, 1:200, Millipore, AB9610), anti-PBXIP1 (Rabbit, 1:100, Abcam, ab84752), anti-PDGFRA (Rabbit, 1:200, Santa Cruz, sc-338), anti-PPP1R17 (Rabbit, 1:200, Atlas Antibodies, HPA047819), anti-SOX9 (Goat, 1:500, R&D Systems, AF3075), anti-SPARCL1 (Goat, 1:300, Novus Biologicals, AF2728), anti-TFAP2C (Rabbit, 1:100, Thermo Fisher Scientific, 14572-1). PBS was used to wash off the primary antibodies, and sections were then incubated with Alexa Fluor secondary antibodies (1:1,000, Life Technologies) for 1 hour at room temperature. Hoechst 33258 was used to visualize the nuclei. Sections were mounted for microscopy with glass coverslips using Aquamount (Thermo Scientific). Images were taken using a Leica TCS SP8 confocal microscope and processed using ImageJ (Fiji). Cortical images spanning from VZ to CP were obtained using a tiling approach in the Leica TCS SP8 and automatically stitched using the Leica software.

## SUPPLEMENTARY FIGURES

**Supplementary Figure S1:**
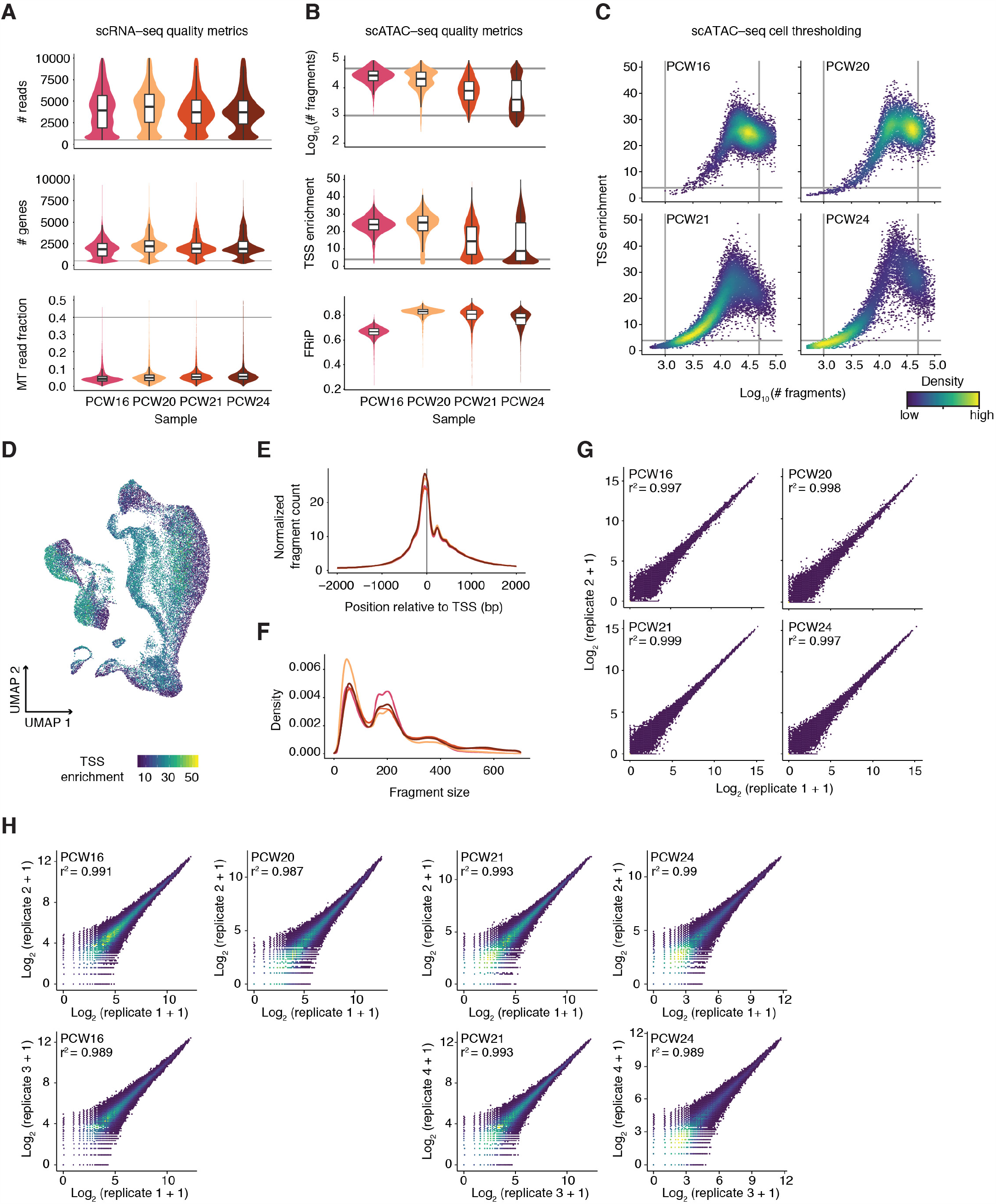
Data quality of scATAC-seq and scRNA-seq libraries. (A) scRNA-seq quality metrics showing the distribution of the number of reads, number of genes, and mitochondrial (MT) gene fraction per cell in each sample. Technical replicates are merged. PCW = postconceptional weeks. (B) scATAC-seq quality metrics showing the distribution of the number of fragments, transcription start site (TSS) enrichment, and fraction of reads in peaks (FRIP) per cell in each sample. (C) scATAC-seq cell thresholding on TSS enrichment and fragment counts. (D) UMAP plot showing the TSS enrichment of each cell. (E) Aggregate normalized fragment count around TSSs for each scATAC-seq sample. (F) Aggregate fragment size distributions for each scATAC-seq sample. (G) Correlation of technical replicates for each scRNA-seq sample. (H) Correlation of technical replicates for each scATAC-seq sample.

**Supplementary Figure S2:**
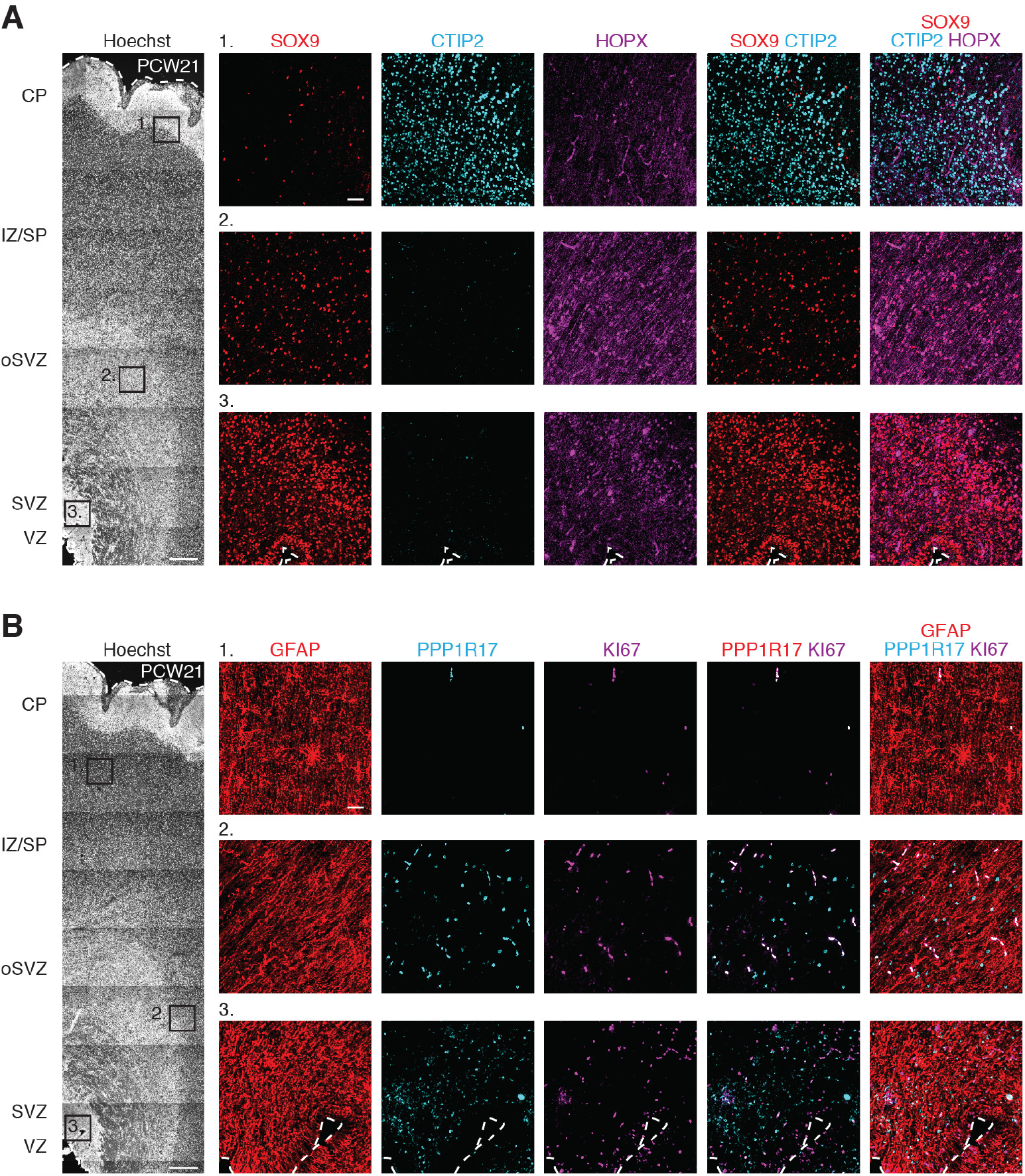
Immunohistochemistry of human cerebral cortex architecture. (A) Immunohistochemistry in PCW21 human fetal cerebral cortex, showing expression of SOX9, CTIP2, and HOPX in the ventricular zone (VZ), subventricular zone (SVZ), outer SVZ (oSVZ), intermediate zone / subplate (IZ/SP), and cortical plate (CP). (B) Immunohistochemistry in PCW21 human fetal cerebral cortex, showing expression of GFAP, PPP1R17, and KI67. Scale bars, 500 μm (A, B), 50 μm (insets A, B).

**Supplementary Figure S3:**
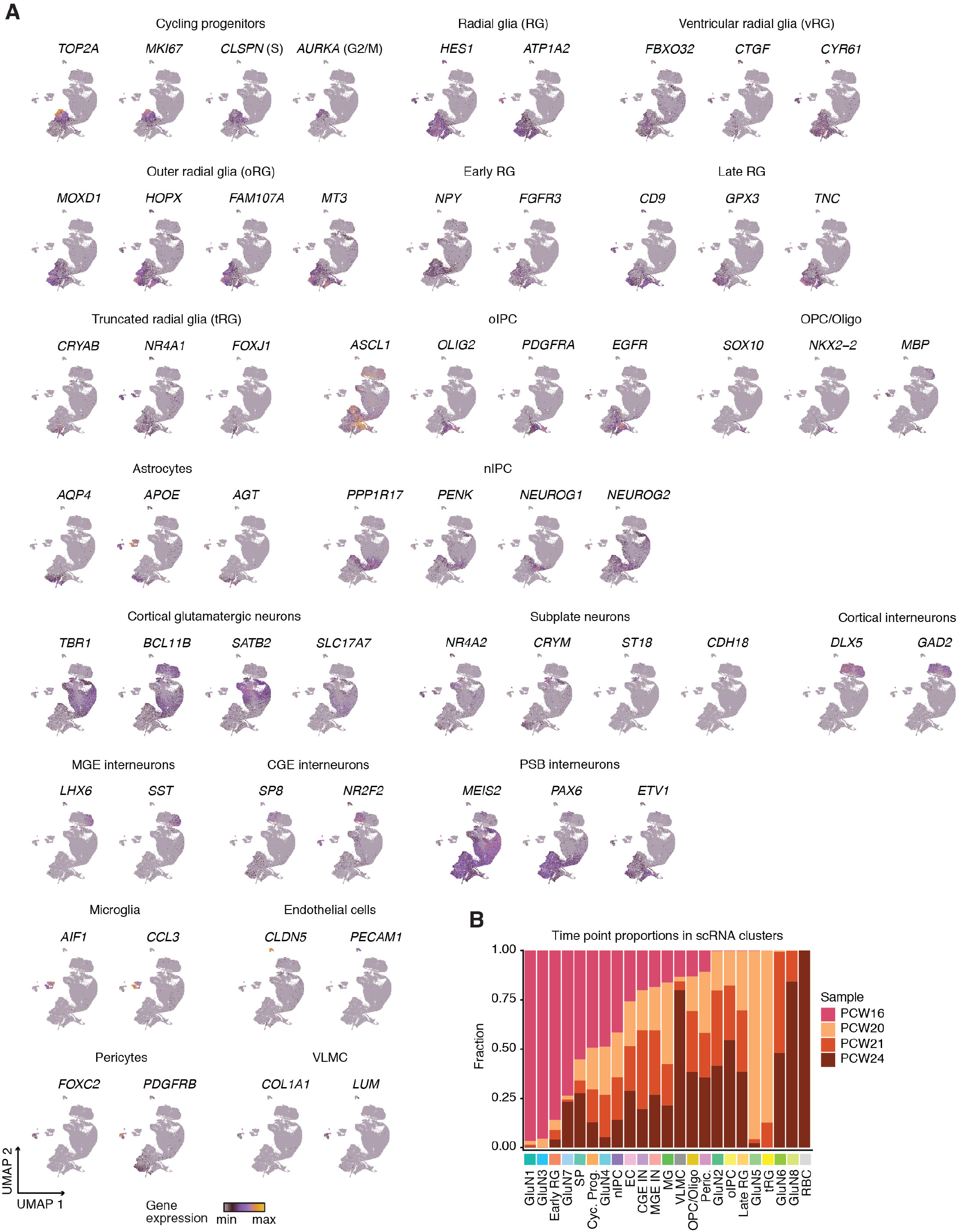
Expression of cell-type specific markers in scRNA-seq data. (A) UMAP plots showing gene expression of cell-type and cluster-specific markers (B) Bar plot showing the sample age composition in each of the scRNA-seq clusters.

**Supplementary Figure S4:**
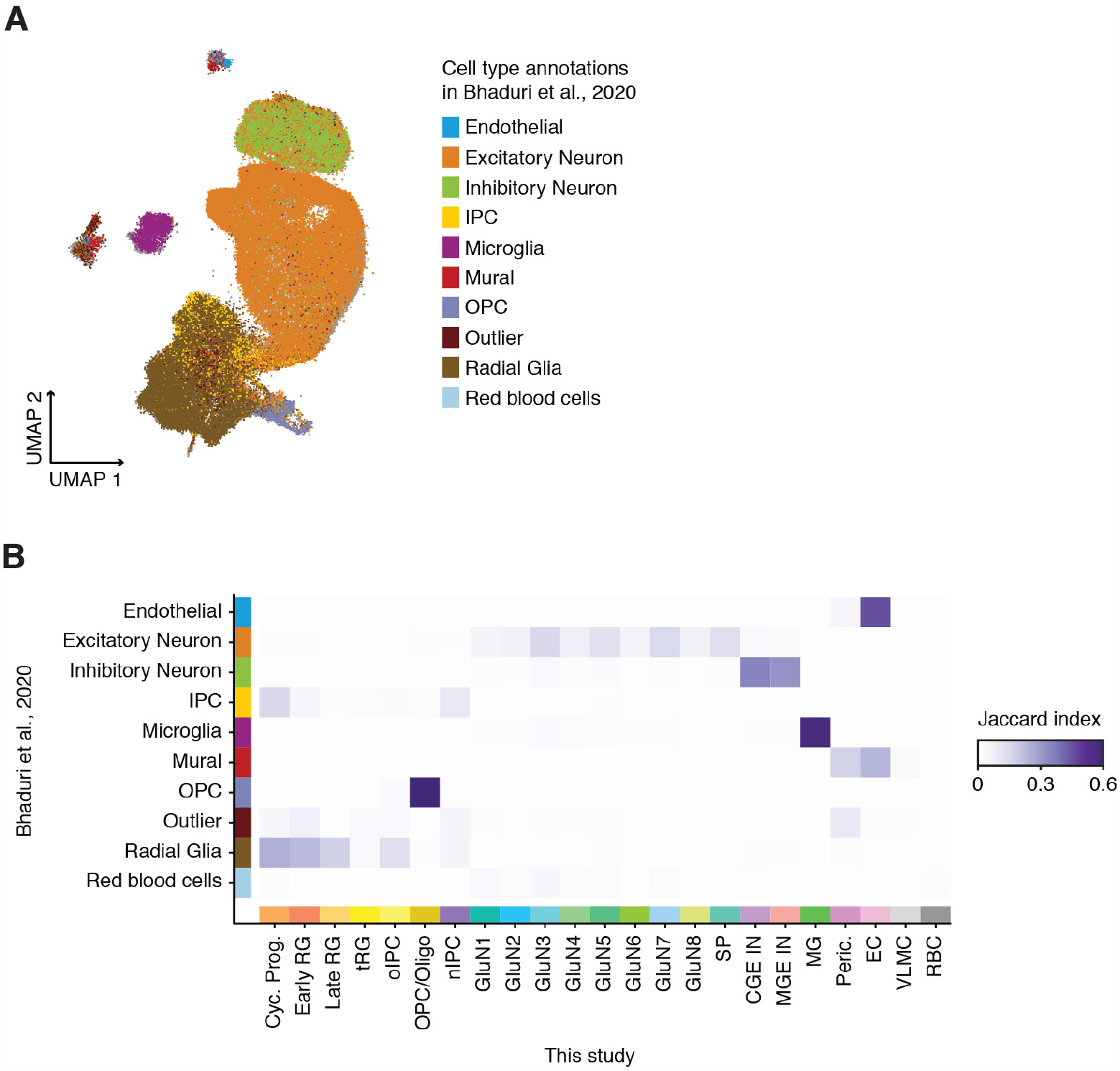
Comparison with external scRNA-seq dataset from human cerebral cortex. (A) Projection of alternate data from Bhaduri et al., 2020 into this scRNA-seq manifold, showing alignment of broad cell types. (B) Jaccard index of genes expressed in clusters from this scRNA-seq dataset and annotated cell types from Bhaduri et al., 2020.

**Supplementary Figure S5:**
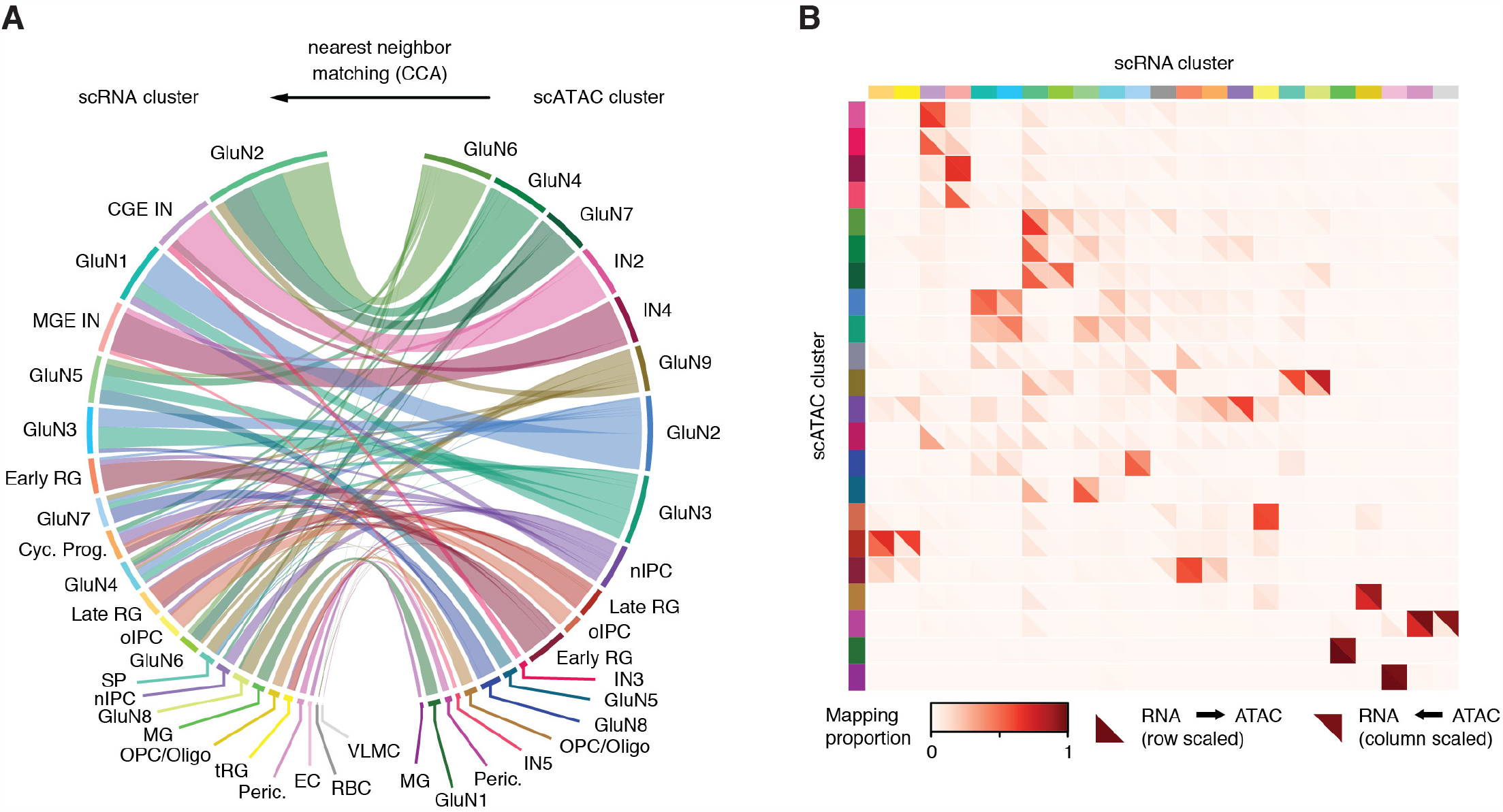
Canonical correlation analysis links scRNA-seq and scATAC-seq datasets in a unified manifold. (A) Ribbon plot showing correspondence of scRNA-seq and scATAC-seq clusters in a shared canonical correlation analysis (CCA) landscape. CCA was derived from expression values in scRNA-seq data matched to gene activity scores from scATAC-seq. (B) Confusion matrix showing the correspondence of cluster annotations across datasets in the CCA. Upper triangles indicate how ATAC clusters match to RNA clusters; lower triangles indicate how RNA clusters match to ATAC clusters. Coloring indicates the proportion of cells mapping for a given pair.

**Figure S6:**
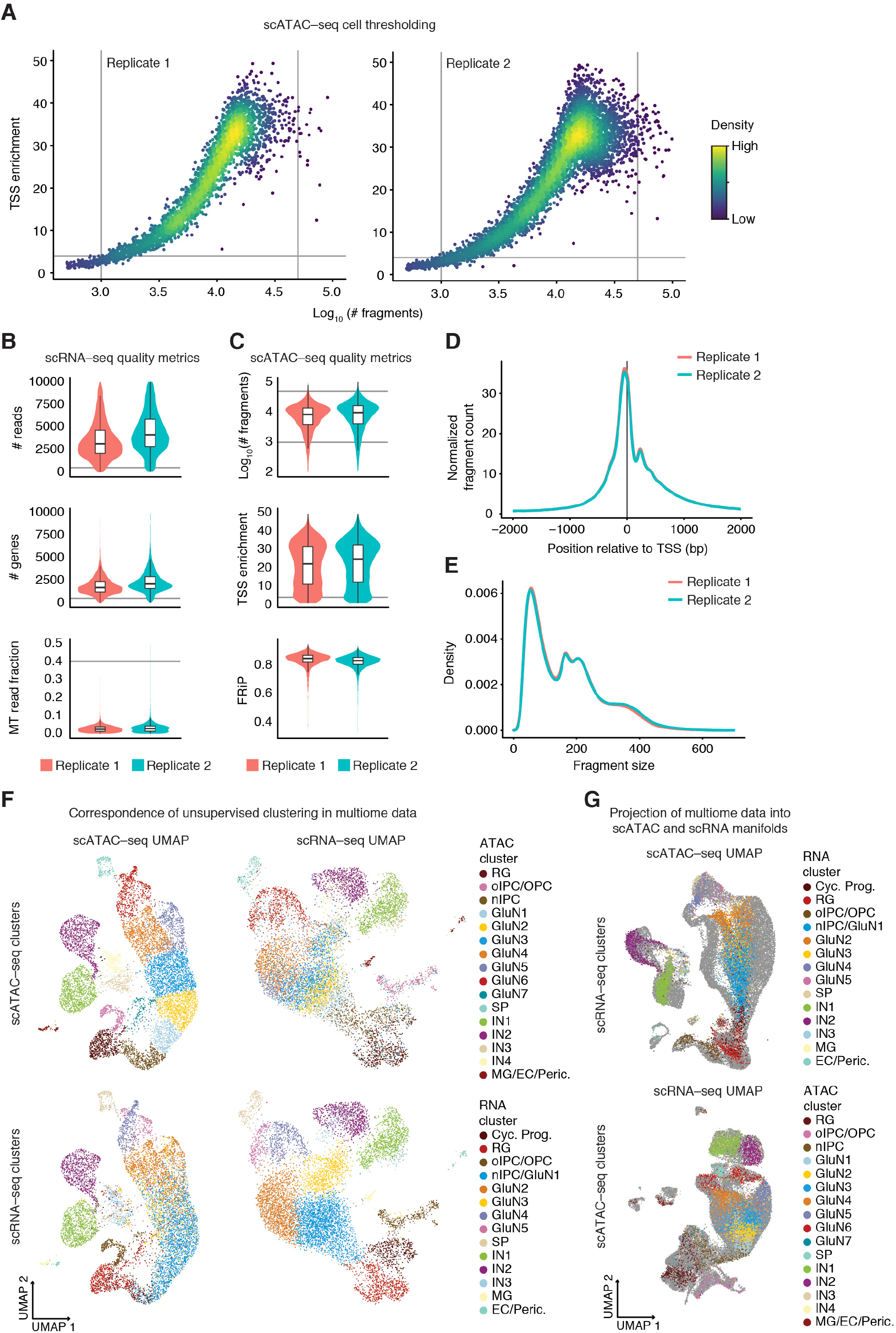
Data quality of scATAC-seq and scRNA-seq multiome data. (A) scATAC-seq cell thresholding on TSS enrichment and fragment counts. (B) scRNA-seq quality metrics showing the distribution of the number of reads, number of genes, and mitochondrial (MT) gene fraction per cell in each biological replicate. Technical replicates are merged. (C) scATAC-seq quality metrics showing the distribution of the number of fragments, transcription start site (TSS) enrichment, and fraction of reads in peaks (FRiP) per cell in each biological replicate. (D) Aggregate normalized fragment count around TSSs for each scATAC-seq biological replicate. (E) Aggregate fragment size distributions for each scATAC-seq biological replicate. (F) UMAP embeddings for multiome scATAC (left panels) and multiome scRNA (right panels). Cells are colored by unsupervised clustering of scATAC counts (top panels) and scRNA data (bottom panels). (G) Projection of multiome scATAC and scRNA data into singleome scATAC (top) and scRNA (bottom) UMAP manifolds.

**Supplementary Figure S7:**
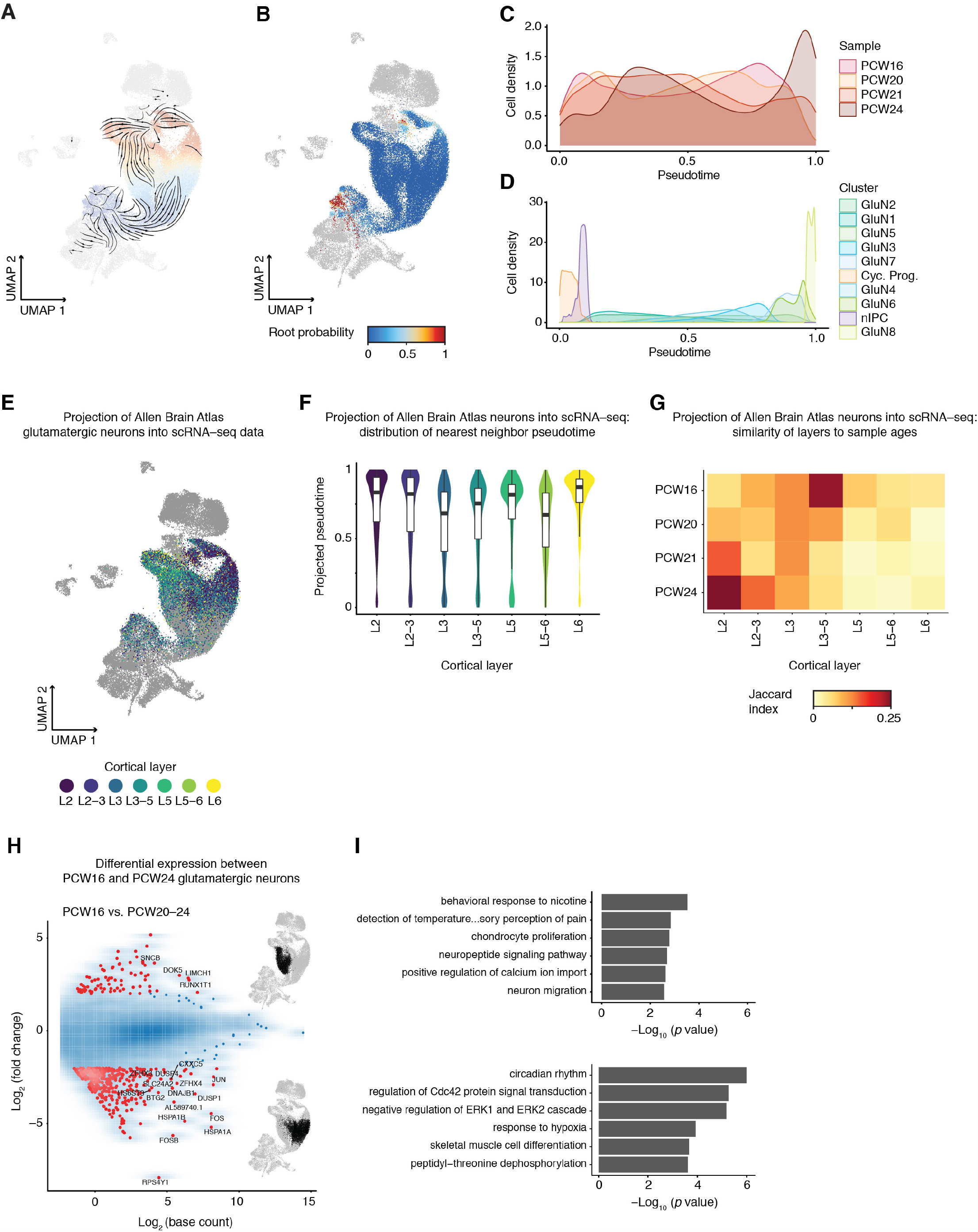
Supplemental analyses to glutamatergic neuron developmental trajectories. (A) RNA velocity streamplot in UMAP space. Aggregate velocities for cells in clusters for glutamatergic neuron trajectories were computed and plotted using scVelo. (B) scVelo root probability in UMAP space. (C) Density plot of sample age for individual cells along the excitatory neuron trajectory pseudotime. (D) Density plot of cell clusters along the excitatory neuron trajectory pseudotime. (E) Projection of adult glutamatergic neurons (Allen Brain Atlas) into scRNA UMAP space. (F) Distribution of excitatory neuron trajectory pseudotime for annotated cortical layers. Fetal cell pseudotime annotation was transferred to adult neurons by nearest-neighbor matching in UMAP space. (G) Correspondence between fetal sample age and annotated adult cortical layers. The heatmap shows Jaccard indices of annotation in adult neurons with fetal gestational age annotation by nearest neighbor matching in UMAP space. (H) MA plot of differential expression between PCW16 and PCW20-24 cells. Genes identified as differentially expressed are shown in red (adjusted p-value < 0.05, |log2(fold-change)| > 2). Cells with 0.2 ≤ annotated pseudotime ≤ 0.8 were compared in PCW16 vs PCW20, PCW21 and PCW24. (I) GO enrichment for genes upregulated (top) and downregulated (bottom) in PCW16 vs PCW20-24 neurons. Enrichments were computed for the gene sets shown in H and the top 6 enrichments are shown for each direction.

**Supplementary Figure S8:**
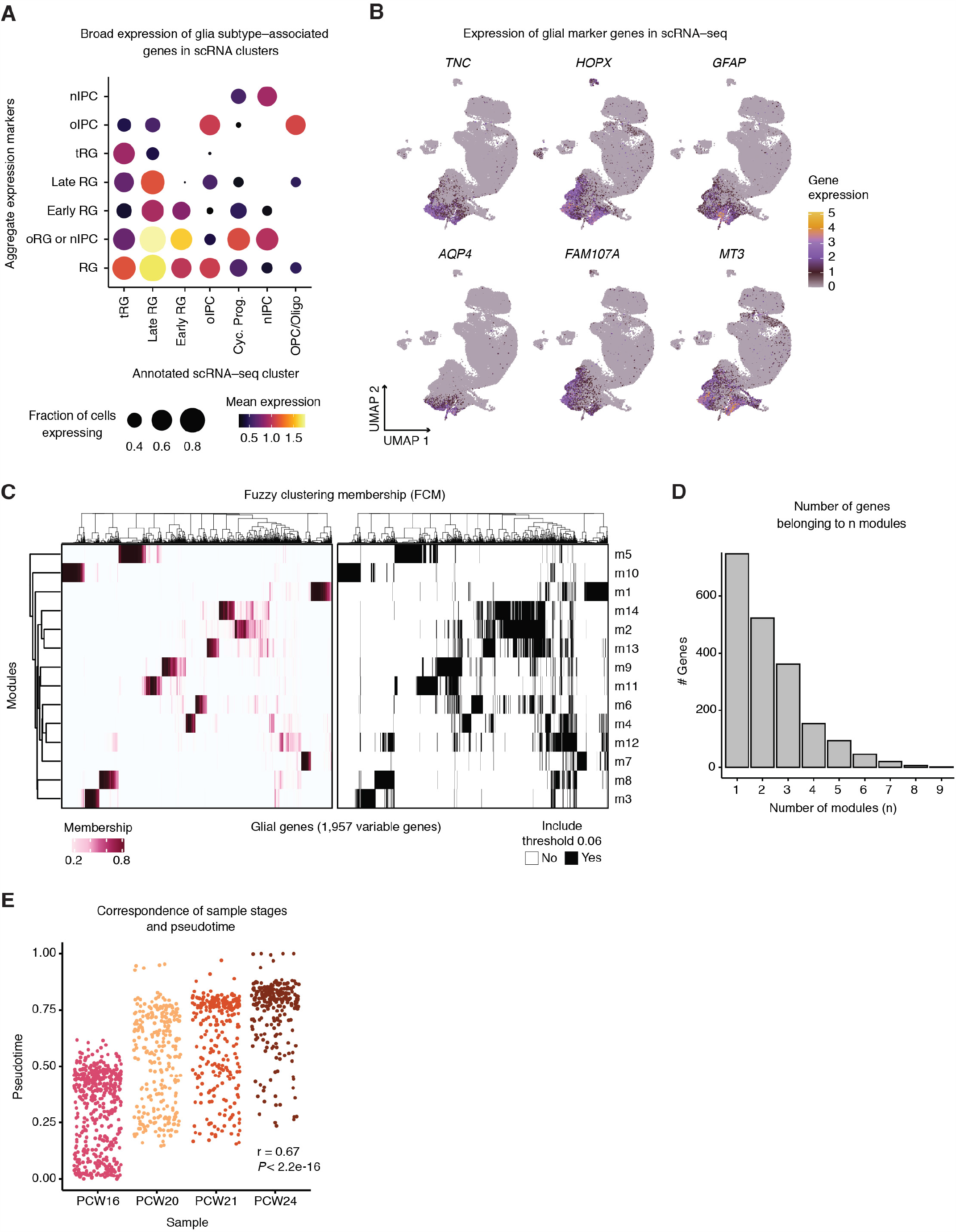
Glial cell characterization using fuzzy c-means clustering. (A) Bubble plot showing gene expression of glial subtype markers in annotated glial clusters. The expression of identifying markers is sometimes evident in several clusters. For each group of markers, the dot size indicates the mean fraction of cells expressing the markers. Color indicates mean expression level. (B) UMAP showing expression of selected glial genes in the scRNA-seq manifold.(C) Membership matrix for fuzzy clustering, showing the fractional membership of each gene (columns) in each module (rows). The right-hand panel shows the memberships, now binarized at a membership threshold of 0.06. (D) Bar plot showing how many genes belong to “n” modules after thresholding. (E) Plot of glial scRNA-seq pseudobulk aggregates. For each aggregate, the sample-of-origin age in postconceptional weeks (PCW) is compared with the pseudotime values (Methods). Pseudotime was strongly correlated with developmental time. Pearson r = 0.67, *P* = 2.2e-16.

**Supplementary Figure S9:**
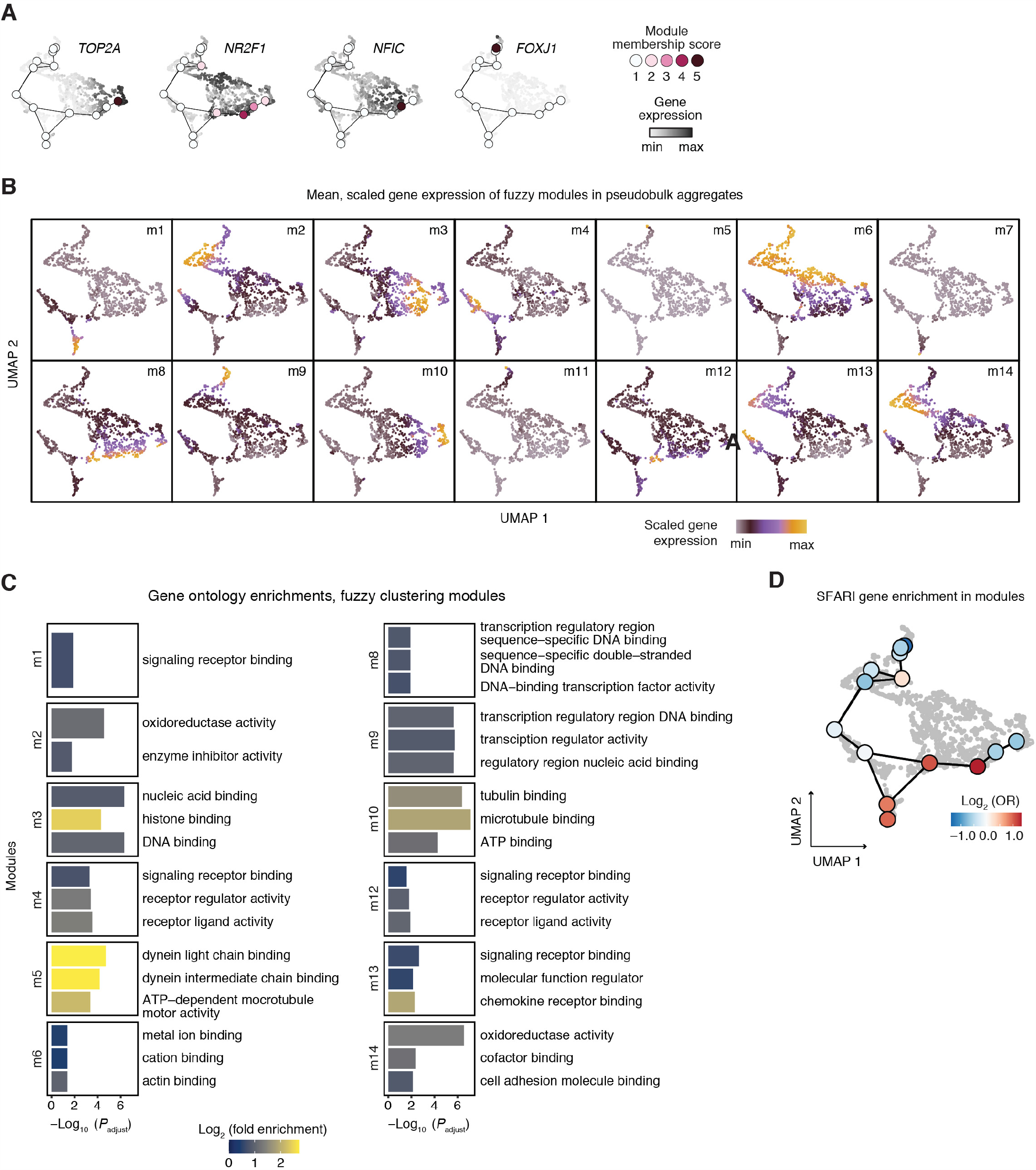
Characterization of fuzzy clusters. (A) Module membership and expression values for genes depicted in **Figure 4C** across pseudotime aggregates. (B) UMAP plots showing the mean, scaled expression of all genes in each module (m1-m14). (C) Gene ontology (GO) enrichments for each module, including the term description. Bar plots represent the −Log_10_ (*P*), with P values adjusted by the Bonferroni method. Bar color indicates the log_2_ fold enrichment for each term. (D) Enrichment of SFARI genes (gene score < 3) in each fuzzy module. Enrichments indicated by color, are shown as the log_2_ odds ratio (OR), and plotted with module centroids in the UMAP of fuzzy clustering cell loadings.

**Supplementary Figure S10:**
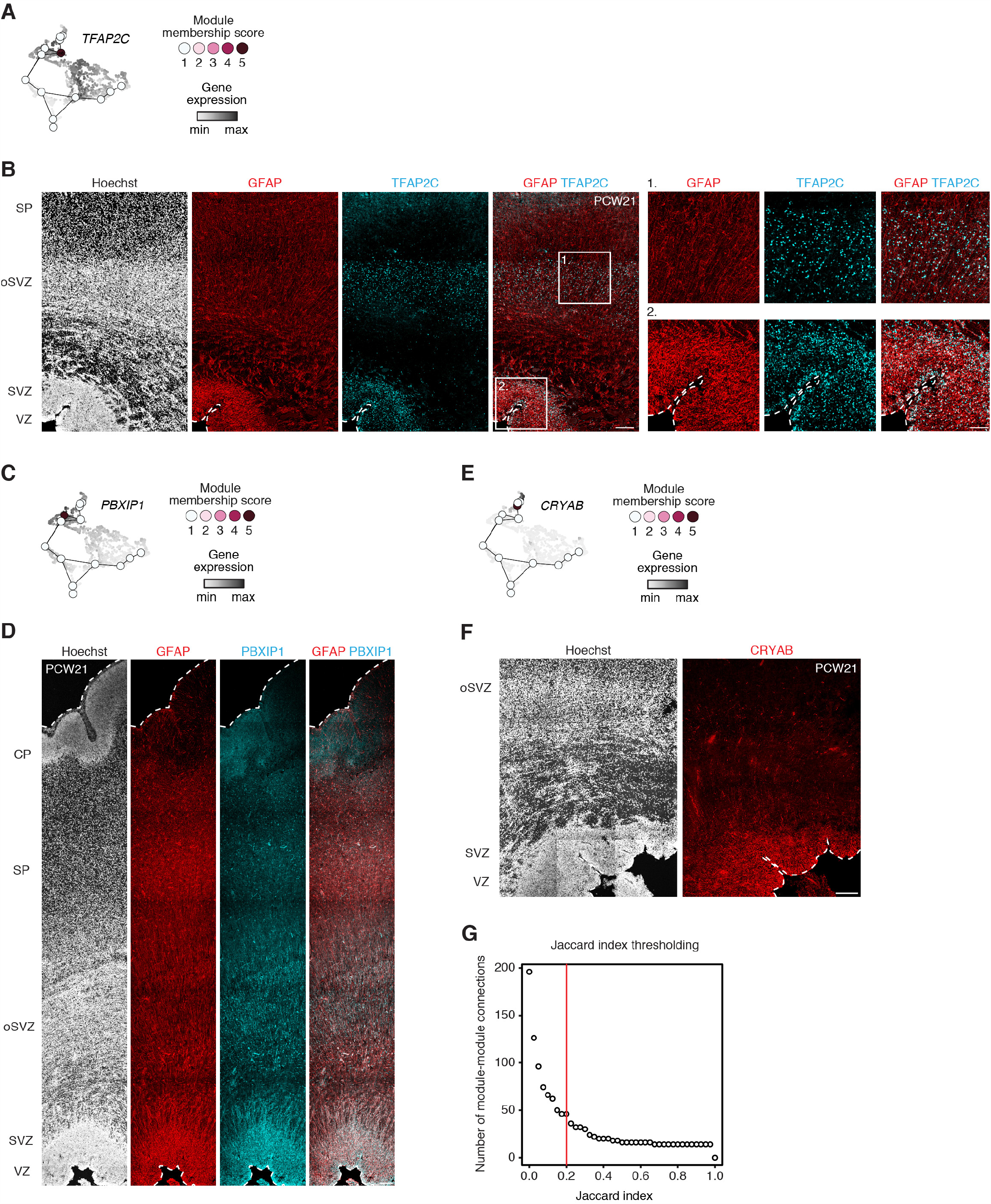
Immunohistochemistry of genes in fuzzy modules. (A) Module membership and expression values for *TFAP2C*. (B) Immunohistochemistry in PCW21 human cerebral cortex showing expression of module m6 transcription factor TFAP2C in the SVZ and oSVZ. (C) Module membership and expression values for *PBXIP1*. (D) Immunohistochemistry in PCW21 human cerebral cortex showing expression of module m2 marker PBXIP1 and colocalization with the astroglia marker GFAP in radial glia in the VZ and oSVZ. (E) Module membership and expression values for *CRYAB*. (F) Immunohistocjemistry in PCW21 human cerebral cortex showing expression of module m9 marker CRYAB in truncated radial glia in the VZ. (G) Plot of the total number of module-module connections at a given Jaccard index threshold. Higher Jaccard thresholds mean fewer connections are “allowed” in the downstream analysis. This plot shows a clear “elbow” behavior at Jaccard > 0.2, which was used to select that threshold. Scale bars, 200 μm (B, D, F), 200 μm (inset B).

**Supplementary Figure S11:**
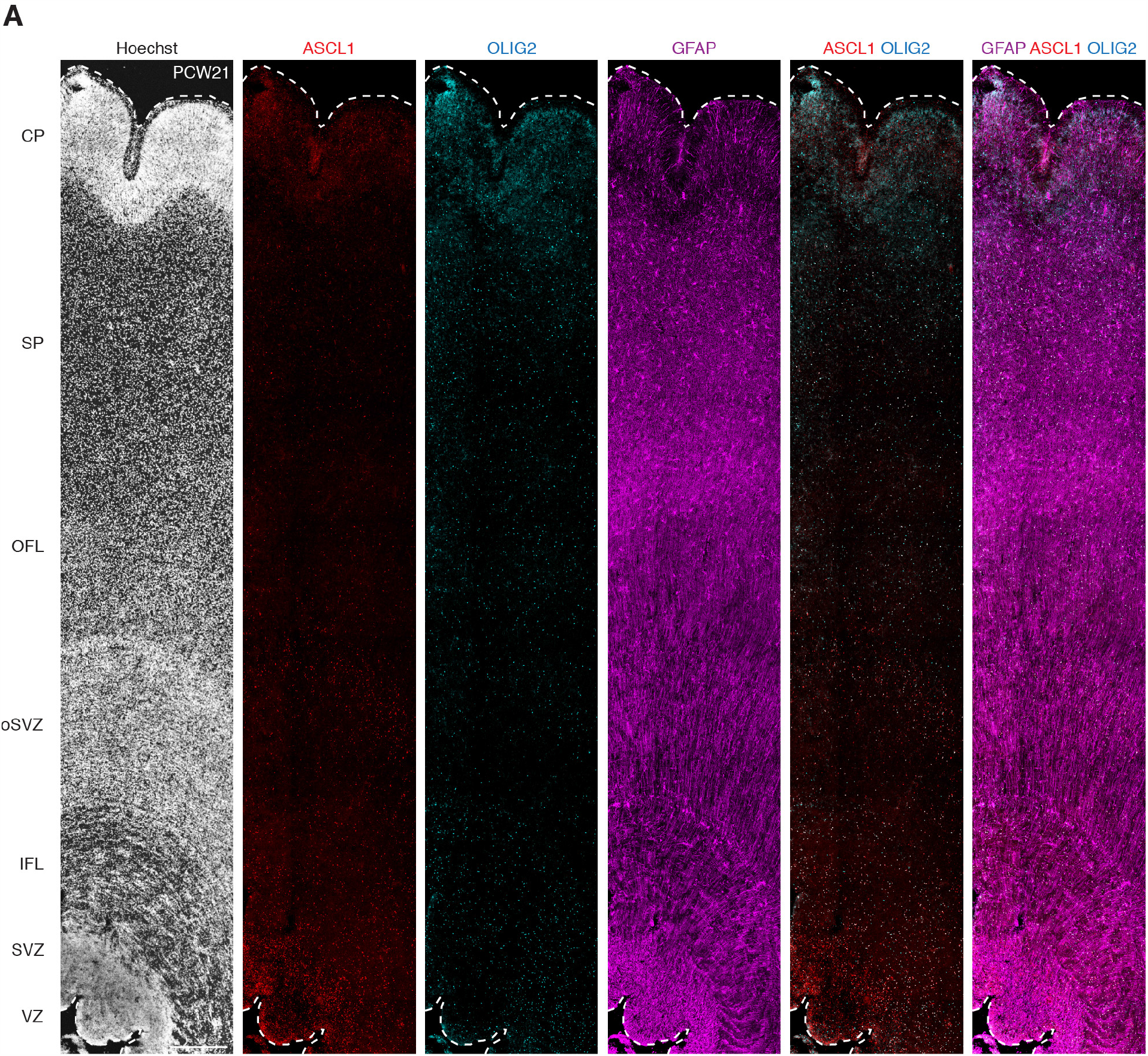
colocalization of OLIG2 and ASCL1 in the human cerebral cortex. (A) Immunohistochemistry in PCW21 human fetal cortex showing expression of ASCL1, OLIG2 and GFAP. ASCL1 and OLIG2 colocalize in the inner and outer fiber layers (IFL, OFL) and SVZ and oSVZ mainly. GFAP shows the radial glial scaffolding. Scale bar, 500 μm (A).

**Supplementary Figure S12:**
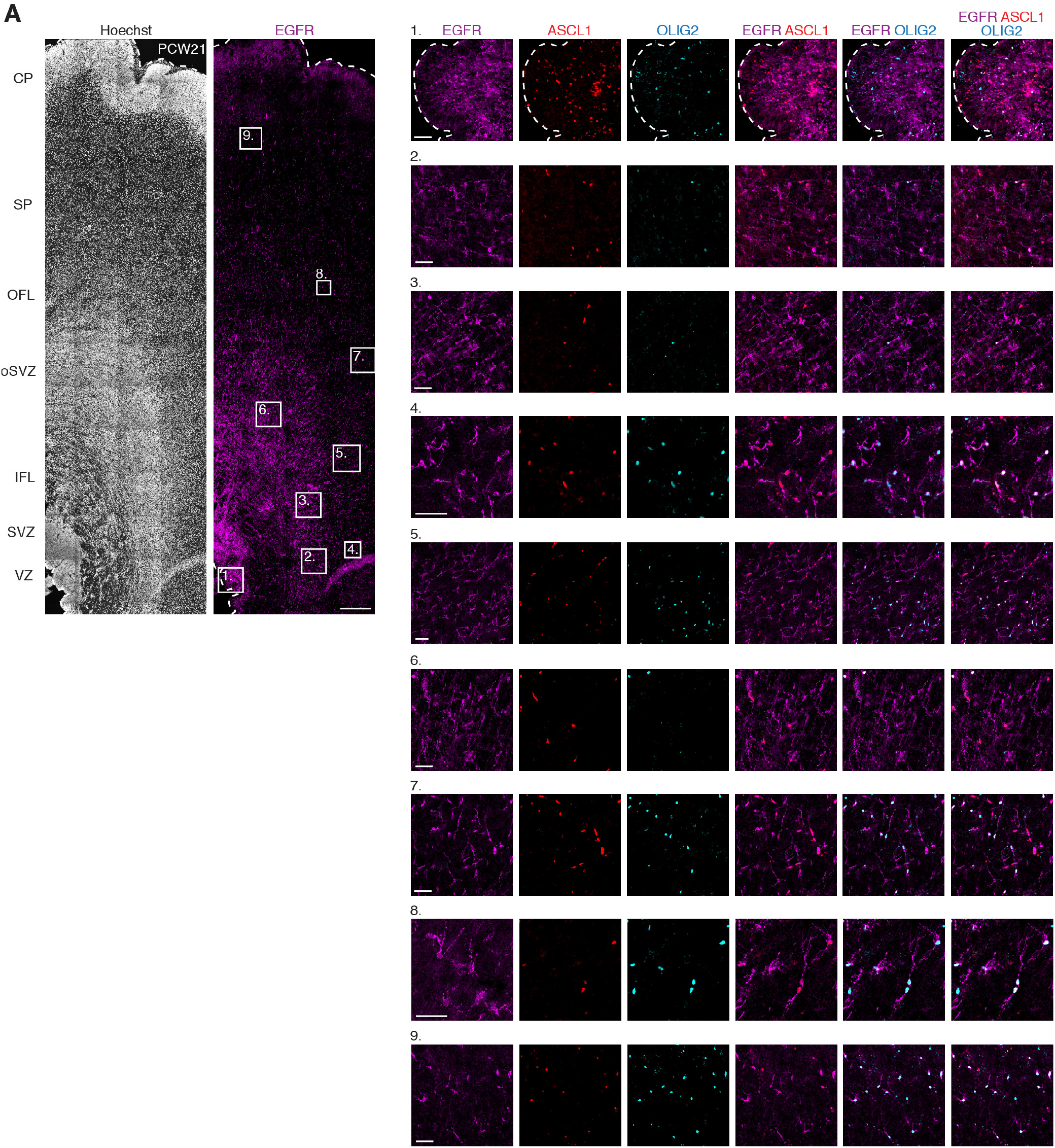
colocalization of OLIG2, ASCL1 and EGFR in the human cerebral cortex. (A) Immunohistochemistry in PCW21 human fetal cortex showing expression and colocalization of modules m1, m4 and m12 genes ASCL1, OLIG2 and EGFR representing oIPCs. Scale bars, 500 μm (A), 50 μm (insets A).

**Supplementary Figure S13:**
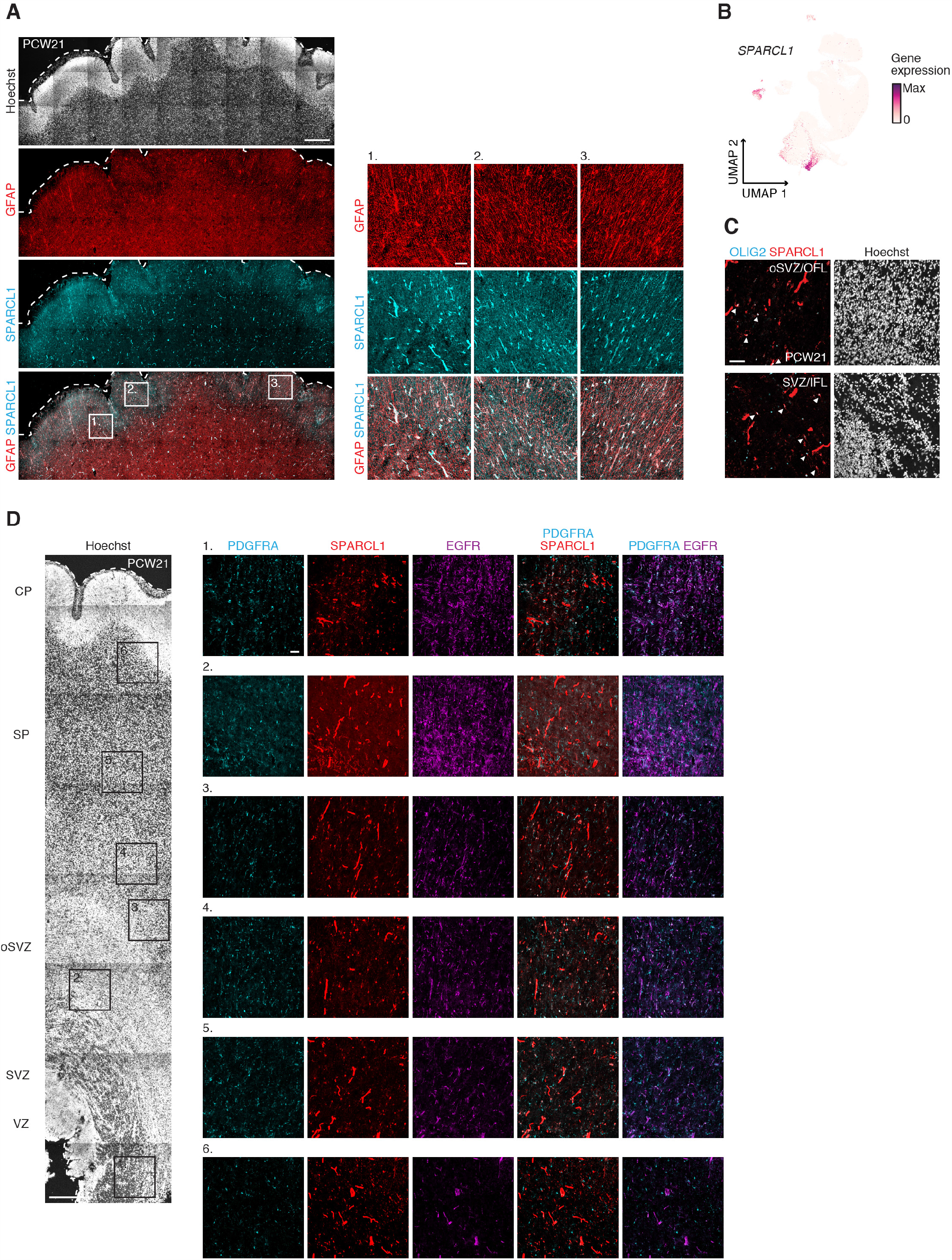
colocalization of astrocyte- and oligodendrocyte-associated markers in the human cerebral cortex. (A) Immunohistochemistry in PCW21 human fetal cortex showing colocalization of the astroglia markers SPARCL1 and GFAP in the cortical plate and subplate. (B) UMAP plot showing *SPARCL1* gene expression. (C) Immunohistochemistry in PCW21 human fetal cortex showing colocalization (white arrowheads) of OLIG2, associated with oligodendrocyte progenitors, and the astrocyte marker SPARCL1 in SVZ/IFL and oSVZ/OFL. (D) Immunohistochemistry in PCW21 human fetal cortex showing colocalization of PDGFRA, SPARCL1 and EGFR. Scale bars, 500 μm (A, D), 50 μm (C, and insets A, D).

**Supplementary Figure S14:**
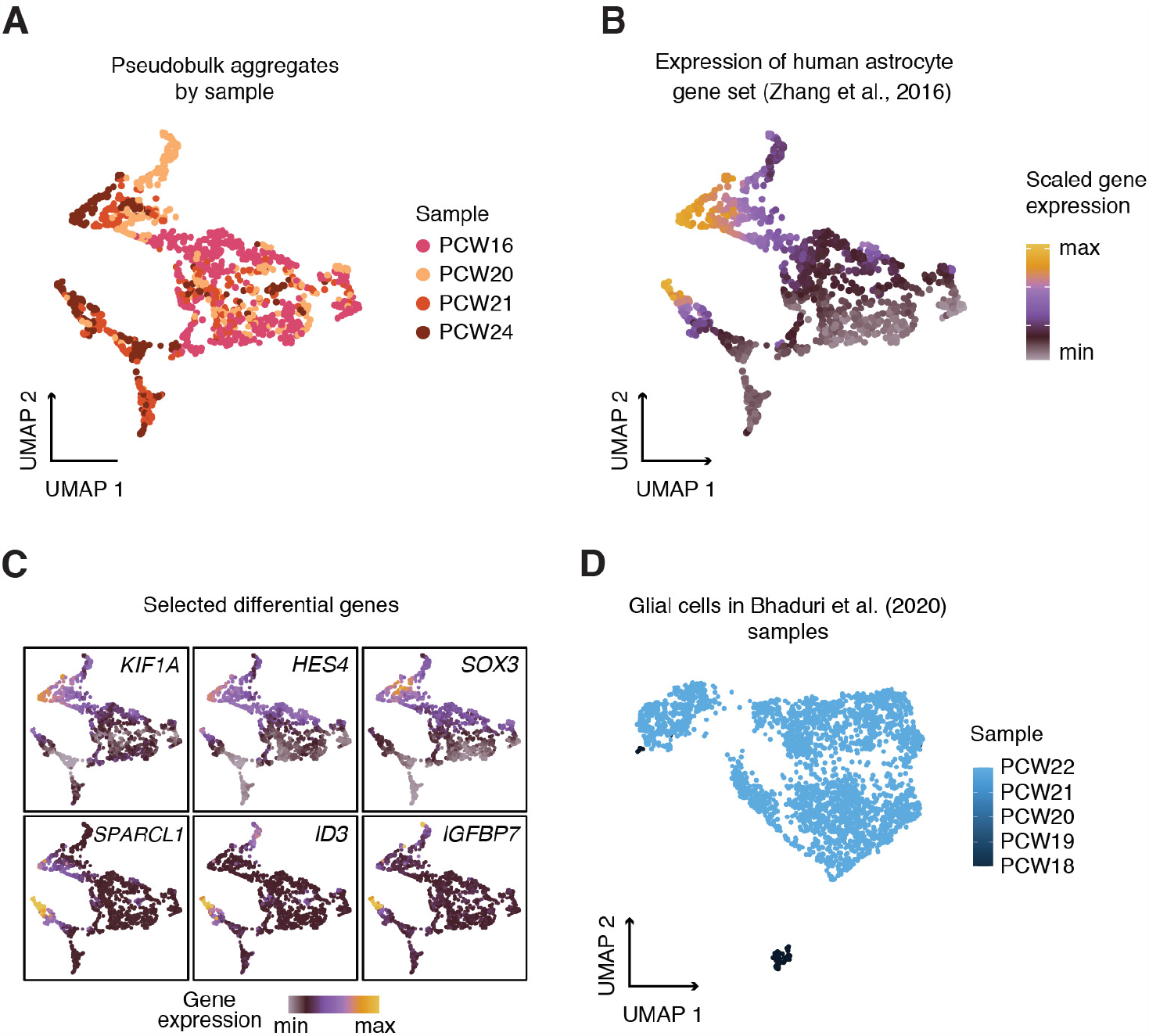
Heterogeneity of astrocyte precursors. (A) Fuzzy clustering-derived UMAP showing pseudobulk aggregates plotted by sample age. (B) Mean scaled expression of human mature astrocyte genes (Zhang et al. 2016) in fuzzy clustering-derived UMAP of scRNA-seq pseudobulk aggregates. (C) Expression of selected differential genes from Figure 4D. (E) UMAP of Bhaduri et al., 2020 fetal astrocyte scRNA-seq dataset, showing sample age.

**Supplementary Figure S15:**
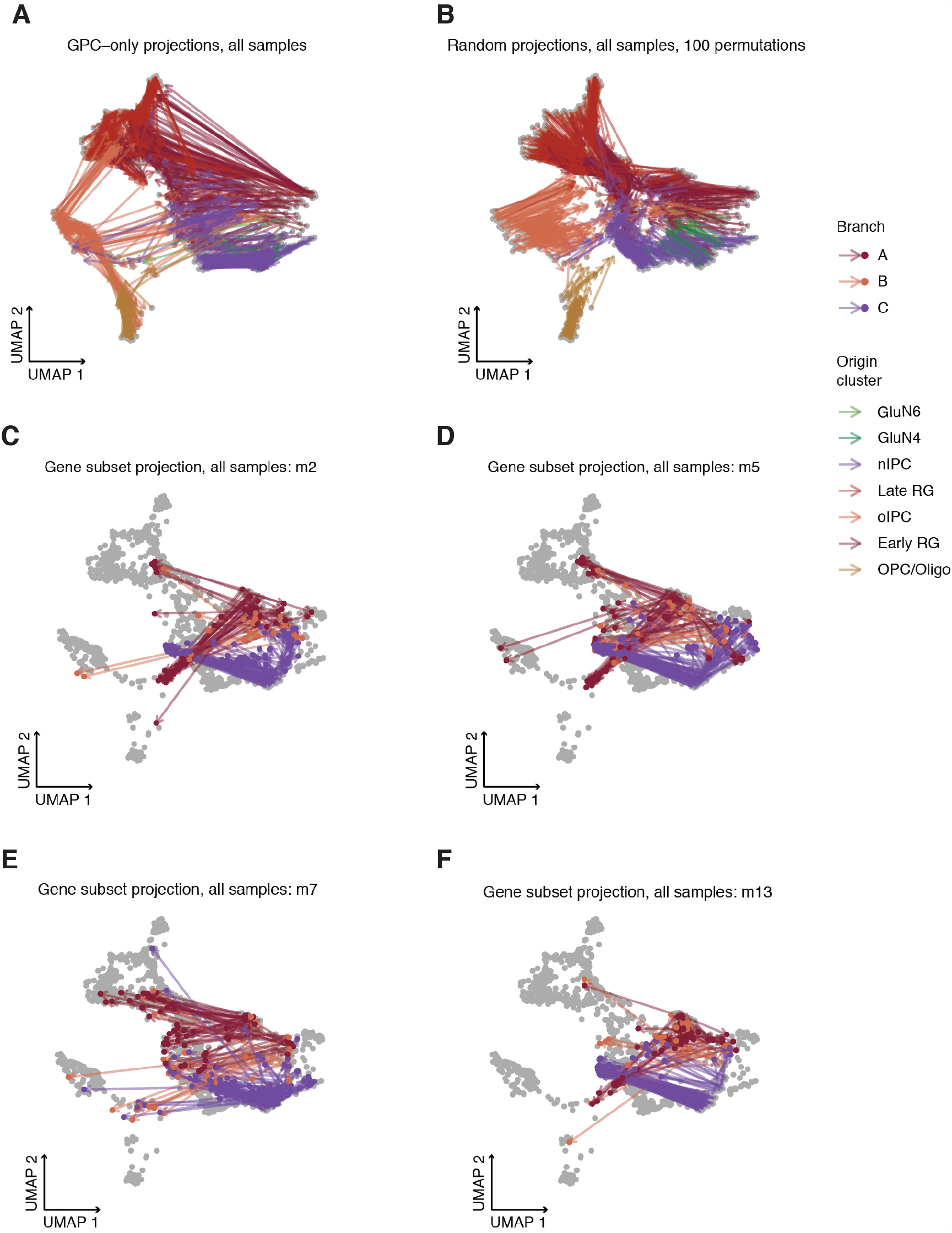
Projection of scATAC-seq aggregates into fuzzy embedding using different gene sets – gene set controls related to GPC analysis. UMAP plots showing the projection of aggregates into the fuzzy clustering-derived low-dimensional embedding. The origin of the arrows represents the original projection coordinates of a particular scATAC-seq aggregate; the arrows point to the new projection coordinates when using only a given subset of genes to make the projection (other genes are imputed as zero-variance features). Colors indicate the scATAC-seq cluster from which the aggregates derive. Panels show projection with only GPC genes (A); random gene sets (B, 100 permuted trials); module m2 genes only (C); module m5 genes (D); module m7 genes (E); module m13 genes (F).

**Supplementary Figure S16:**
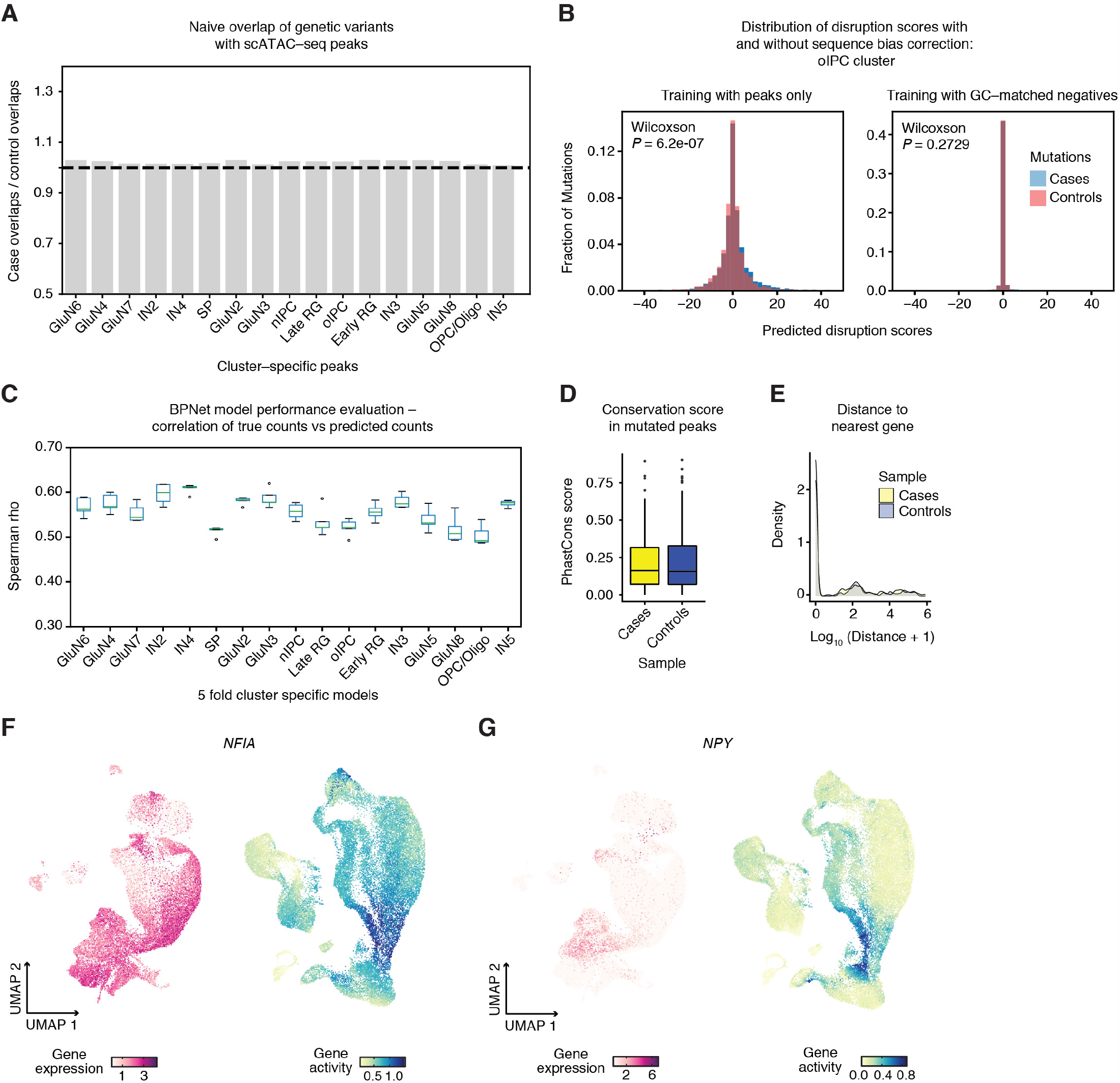
Supplementary characterization of BPNet model performance and mutation vignettes. (A) Enrichment of cases versus control mutations using naïve overlap with cluster-specific ATAC-seq peaks, showing rele vance of the deep learning model to capture pathogenic disruptions. (B) Distribution of disruption scores for case and control mutations using different training paradigms. Data are shown for the oIPC cluster. On the left, using only scATAC-seq peaks as the basis for training, there is a systematic difference between cases and controls (Wilcoxson test *P* = 6.2e-7). On the right, when training is given GC-matched negatives, disruption scores are substantially more conservative, and the distributions are matched (*P* = 0.27). (C) Performance evaluation of BPNet cluster-specific models, computed by calculating the rank correlation between true counts in the cluster and predicted counts. Data are from 5-fold cross-validated training. (D) Conservation scores in cases versus controls, showing that trivial genomics metrics do not explain the observed prioritized mutations. (E) Distance to the nearest gene in cases versus controls, showing that trivial genomics metrics do not explain the observed prioritized mutations. (F) UMAP plots of gene expression (magenta) and gene activity (viridis) for *NFIA*. (G) UMAP plots of gene expression (magenta) and gene activity (viridis) for *NPY*.

